# *In Vivo* Selection of anti-glioblastoma DNA aptamer-drug conjugates in an orthotopic patient-derived xenograft model

**DOI:** 10.64898/2026.02.16.706148

**Authors:** Caroline D. Doherty, Sonia Jain, Lauren L. Ott, Katie K. Bakken, Brandon A. Wilbanks, Danielle M. Burgenske, Keenan S Pearson, Jessica I. Griffith, Zichen Tian, Jeffrey A. Meridew, Saigopalakrishna S. Yerneni, William F. Elmquist, Jann N. Sarkaria, L. James Maher

**Affiliations:** Department of Molecular Pharmacology and Experimental Therapeutics, Mayo Clinic Graduate School of Biomedical Sciences, Rochester, MN 55905; Department of Biochemistry and Molecular Biology, Mayo Clinic Graduate School of Biomedical Sciences, Rochester, MN 55905; Department of Radiation Oncology, Mayo Clinic, Rochester, MN 55905; Department of Pharmaceutics, University of Minnesota; Department of Laboratory Medicine and Pathology, Mayo Clinic, Rochester, MN 55905

**Keywords:** *in vivo* SELEX, conjugate SELEX, aptamer, aptamer drug conjugate, ApDC, MMAE, neuro-oncology, glioblastoma, GBM, PDX model, brain tumor, antibody drug conjugate, ADC

## Abstract

Glioblastoma (GBM) is an aggressive, high-grade glioma with a near-universally fatal prognosis. Therapeutic failure is often attributed to the highly selective blood brain barrier (BBB), the diffuse infiltrative nature of the tumor, and the marked intratumoral heterogeneity of GBM. Although antibody drug conjugates ADCs have shown promise for high grade gliomas such as GBM, efficacy is limited by ADC size. Aptamers—short, synthetic, single-stranded DNA or RNA molecules—can be ∼6-fold lower in molecular weight than IgG antibodies and have the potential to cross the intact BBB. Unlike other nucleic acid-based therapies, aptamer function arises from three-dimensional shape rather than genetic coding. Here we aim to replace the targeting component of the ADC paradigm with a DNA aptamer, thus creating an aptamer-drug conjugate (ApDC). We employed *in vivo* SELEX using an orthotopic patient-derived xenograft (PDX) GBM mouse model and a vast (∼100 trillion 80-mer sequences) ApDC library. We report the results from this first in vivo ApDC selection of its kind. We characterize target tissue binding *ex vivo*, cell association, biodistribution, and pharmacokinetics from this selection. This study exemplifies an unbiased approach to a problem that rational design has yet to overcome, offering a new direction for GBM therapeutic development.

## Introduction

Antibody drug conjugates (ADCs) have recently made favorable inroads in oncology treatment. At the beginning of 2019, the FDA had approved only four ADCs for cancer treatment; as of 2026 this number has risen to 15 (1, 2). ADCs are comprised of three components: (1) a targeting antibody, (2) a cleavable or non-cleavable linker, and (3) a highly cytotoxic drug. ADCs often are touted as “biological guided missiles,” embodying Paul Ehrlich’s concept of “magic bullets” (1, 3, 4). Despite the successes of ADCs in hematological malignancies and some solid tumors, similar efficacy has not yet been achieved in high grade brain tumors (5–8).

Glioblastoma (GBM) is the most common high grade brain tumor in adults. It is highly aggressive with a median survival time of ∼14 months (9). Treating GBM is a therapeutic challenge in part due to the blood brain barrier (BBB), which prevents many therapies from reaching the tumor from the peripheral circulation (10). Additionally, GBMs exhibit diffuse, infiltrative growth patterns and display high intratumoral heterogeneity, both of which confer drug resistance and tumor recurrence (10). ADCs have shown promise for GBM, but due to the relatively large size of the antibody targeting domain, their efficacy is dependent on BBB leakiness, a parameter that is heterogenous across patients and within tumors (6). Efforts to exploit receptor-mediated pathways, such as receptors for epidermal growth factor (EGF) or transferrin, have been pursued. However, results suggest that high-affinity binding promotes robust BBB endothelial cell uptake and poor transcytosis into the brain parenchyma (11, 12). More importantly, to date, all clinical trials evaluating anti-EGFR and anti-transferrin ADCs for GBM have been terminated due to toxicity or lack of efficacy (8, 13). Furthermore, antibodies, similar to small molecules, are based on rational target selection based on a single tumor antigen. Thus, antitumor activity is inherently constrained, especially in highly heterogenous tumors like GBM, which have propensity for antigen escape as exemplified by GBM clinical trials for EGFR-targeting CAR T-cell therapy (14, 15).

Aptamers are small, synthetic RNA or DNA molecules that fold into unique three-dimensional shapes and act as antibody analogs, yet are typically ∼6-times lower in molecular weight than an IgG antibody (10). Unlike antisense and anti-gene nucleic acids, aptamers function as three-dimensional shapes, unrelated to genetic coding. Like antibodies, aptamer structure serves as the basis for highly-specific binding: aptamers can bind ligands with equilibrium dissociation constants in the sub-nM range, similar to antibodies (16, 17). However, aptamers are typically nonimmunogenic and nontoxic, even at doses 1000-fold higher than typical antibody therapies (18). Rather than being engineered to bind to specific targets, aptamers can be selected through an agnostic process termed <u>S</u>ystematic <u>E</u>volution of <u>L</u>igands by <u>Ex</u>ponential enrichment (SELEX). SELEX is characterized by repetition of rounds based on three steps: (1) incubation of a random library of ∼100 trillion unique molecules (10^14^) with a simple or complex target of interest, (2) removal of all unbound molecules, and (3) rewarding target-bound molecules with PCR amplification.

A fundamental advantage of SELEX is that selection can proceed without a predefined molecular target. At the final round, the library is typically enriched for multiple “winning” molecules, and in selections against complex tissues or tumors, a cocktail of these dominant aptamers will likely target multiple tumor antigens. In the case of *in vivo* SELEX, molecules are also rewarded for homing to the target tissue—using whatever unspecified means is available to them. Thus, the ‘fittest’ targeting strategy, typically unknown, will be employed by winning molecules. Until recently, selection of aptamers relevant to neuro-oncology has been limited primarily to cell-SELEX (10). The challenge with this method is that the resulting aptamers, like antibodies, may bind a target protein or cell with high affinity, but their biodistribution, pharmacokinetics (PK), and target homing *in vivo* are not optimized by the selection. Consequently, high-affinity binding alone may be insufficient to achieve therapeutic efficacy *in vivo*. We recently completed the first successful *in vivo* SELEX in an orthotopic patient-derived xenograft (PDX) GBM mouse model (19). The *in vivo-*selected DNA molecules demonstrated impressive biodistributions and target cell binding (19). However, when these molecules were subsequently modified by conjugation to a large ADC linker and cytotoxic cargo, they lost their ability to bind target cells in culture (Supplemental Figure S1). We hypothesized that incorporating drug conjugation into the selection process would facilitate the identification of aptamers optimized for conjugate function *in vivo*.

Given the limitations of ADCs, the therapeutic challenges associated with GBM, and the unique benefits of aptamers, we further hypothesized that using *in vivo* SELEX with an ApDC library would allow replacement of the antibody domain of an ADC with *in vivo* trained ApDCs, thus improving the biodistribution, pharmacokinetics, and the number of tumor antigens targeted simultaneously. In this work, we build from ADC technology by utilizing a common ADC linker and toxin, monomethyl auristatin E (MMAE)-p-aminobenzylcarbamate (PAB)-valine-citrulline (VC)-PEG_4_. We first show that our choice of the common ADC cytotoxic anti-mitotic agent, MMAE, as a generic cytotoxic placeholder, is compatible with PCR and SELEX conditions required for *in vivo* selection. We then demonstrate successful *in vivo* SELEX with an ApDC library conjugated to MMAE. We show that the resulting molecules bind the target tumor tissue and cells more strongly than an untrained ApDC and demonstrate encouraging biodistributions and pharmacokinetics in live mice. Leveraging our previous work using a nearly identical *in vivo* SELEX strategy, we note striking differences in results for MMAE conjugates.

The present study focuses on *in vivo* selection methodology and results for ApDCs using MMAE conjugation as an example for methods development. Homing and specificity characteristics of the selected ApDCs are reported in detail. Future studies will address the GBM tumor toxicity of the selected reagents.

## Materials and Methods

### MMAE-conjugated DNA primers

Trans-cyclooctene-tetrazine (TCO-Tz) click chemistry was used to synthesize the initial generation of 5’MMAE-DNA-primers used in Figure 1 (Supplemental Figure S2A). Following these preliminary tests, more consistent results were obtained by commercial purchase of 5’MMAE-conjugated primers due to unpredictable batch-to-batch differences in reactivity of 5’TCO primers (Supplemental Figure S2A-B). For preliminary tests, MMAE-Tz was obtained from Levena Biopharma (custom synthesis) and reconstituted in batches at 10 mM in DMSO (Sigma Aldrich #276855). 5’TCO-DNA-primers were later obtained from Biosynthesis and were reconstituted with water at 100 µM concentration. To conjugate, MMAE-Tz solution was added to 5’TCO-oligonucleotide at a volume ratio of 1 μL for every 10 μL of oligonucleotide. The reaction was mixed by pipetting and allowed to stand at room temperature for at least 1 h.

**Figure 1.**
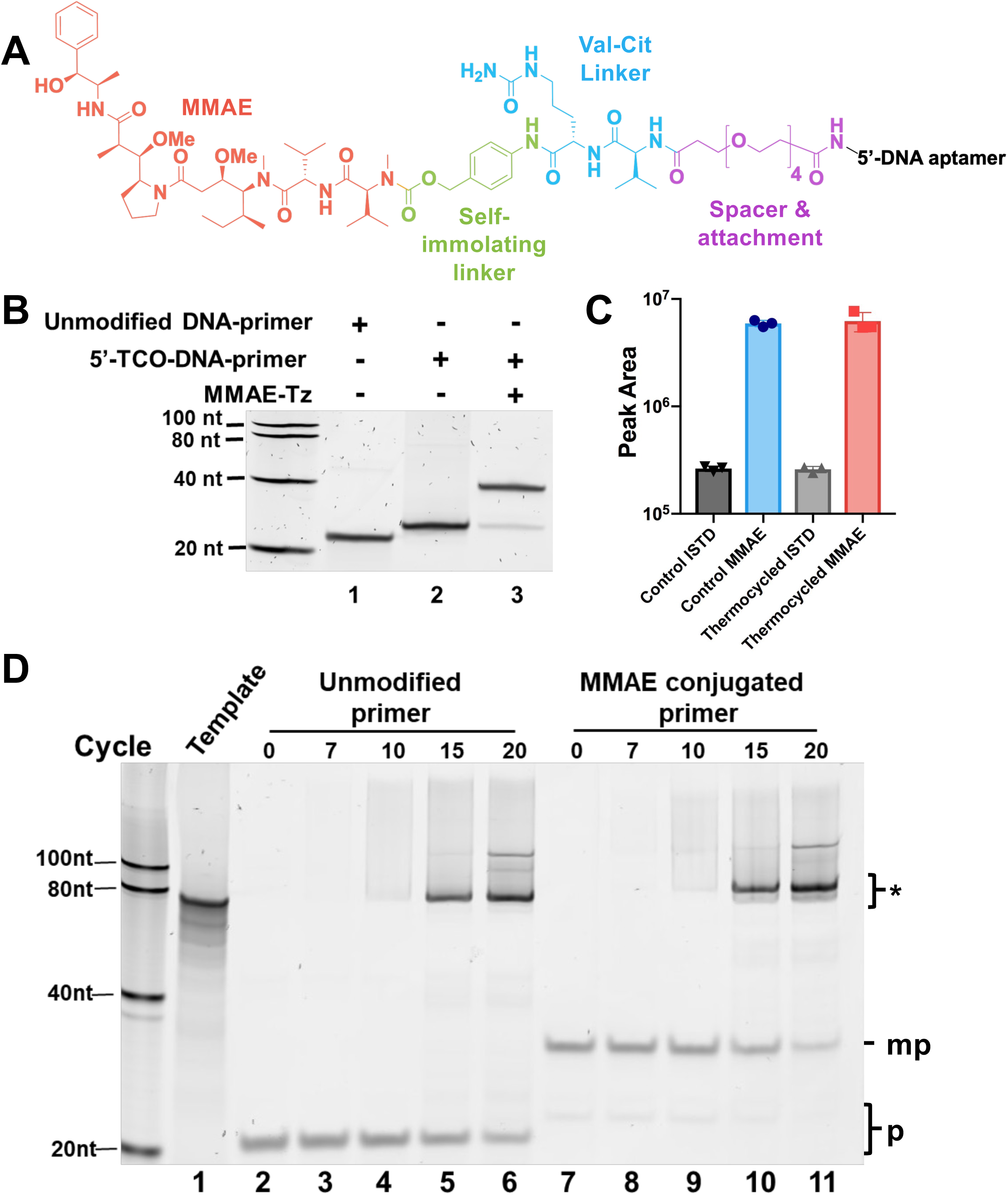
Development of ApDC libraries with cargo compatible with SELEX. A. Chemical structure of PEG4-Val-Cit-PAB-MMAE conjugated to the 5’ end of a DNA aptamer. B. A polyacrylamide denaturing gel post-stained with SYBR gold showing that MMAE-PAB-Val-Cit-PEG4 can be conjugated to DNA primers by trans-cyclooctene-tetrazine (TCO-Tz) click chemistry. C. Confirmation of MMAE structural stability by LC-MS following 20 rounds of thermal cycling conditions. ISTD= internal standard D. Confirmation that Taq polymerase could incorporate a 5’MMAE conjugated DNA primer. Two PCR reactions were run: one with an unmodified forward primer (lanes 2-6) and one with 5’MMAE conjugated forward primer (lanes 7-11). Samples were taken throughout the PCR reaction and run on a native polyacrylamide gel and post-stained with SYBR gold. p=primers, mp = 5’MMAE conjugated primer, and *=amplicon

Subsequent to experiments documented in Figure 1, 5’MMAE primers were purchased from Integrated DNA Technologies based on conventional conjugation of 5’-amino-modified oligonucleotide primers with NHS-ester-Val-Cit-PAB-MMAE (BroadPharm #BP-41657 or ThermoFisher #NC3792832) and purification of the product by HPLC.

### Conjugation confirmation

To confirm MMAE conjugation to the DNA primer, 1 pmol of material was mixed with formamide and heated at 95°C for 5 min. The DNA was subjected to electrophoresis through a 16% denaturing polyacrylamide gel [7.5 M urea, 19:1 acrylamide : bisacrylamide (Bio-Rad #1610144)] for 45 min. The gel was stained with SYBR gold (Invitrogen #S11494) and imaged with a Typhoon imager (Amersham).

### MMAE thermostability

To confirm the chemical stability of MMAE to thermal cycling consistent with PCR, free MMAE was subjected to 20 thermal cycles (1 min at 95°C; 15 s at 94°C, 35 s at 50°C, 30 s at 72°C; hold at 4°C). Samples before and after thermal cycling were analyzed by LC-MS/MS.

### Biological activity of MMAE subjected to thermal cycling

To evaluate the cytotoxicity of free MMAE following thermal cycling, MMAE was diluted in water to 10 µM and was subjected to 25 thermal cycles: 95°C, 60 s; 25× (94°C, 15 s; 50°C, 35 s; and 72°C, 30 s); 4°C. The sample was then heated for a further 5 min at 90°C and placed on ice for 10 min to simulate the snap cool protocol applied during preparation of ApDCs. A CellTiter-Glo assay (Promega #G7571) was used to determine aptamer toxicity. G43-eGFP cells were seeded at 2, 500 cells/well in 50 µL complete media. Free untreated and thermal cycled MMAE were diluted to 10× concentrations in PBS containing 1 mM MgCl_2_. Complete media (40 µL) spiked with 0.1 mg/mL sheared salmon sperm DNA was added to the cells followed by 10 µL of the MMAE solutions. Six d post-treatment, a CellTiter-Glo assay was performed according to the manufacturer’s protocol.

### PCR incorporation of 5’MMAE-conjugated primers

Two 100 µL PCR reactions were assembled with SYBR green qPCR mix (55 µL QuantaBio PerfeCTa #95071-012, 11 µL 20-nt forward primer (5 µM) or MMAE-conjugated forward primer (5 µM), 11 µL 20-nt reverse primer (5 µM), 20 µL water, and 7.5 µL 80-nt template DNA (1 nM). The 5’MMAE-conjugated primer was synthesized using TCO-Tz click chemistry. Reactions were each divided into five 20 µL aliquots. Aliquots (20 µL) were removed after 0, 7, 10, 15, or 20 PCR cycles and subjected to electrophoresis through a 10% native polyacrylamide gel. The gel was first imaged for SYBR green to confirm the presence of duplex amplicons and then stained with SYBR gold and imaged on a Typhoon imager.

### Random DNA library synthesis for *in vivo* SELEX

DNA oligonucleotides (Supplemental Table S1) were obtained from Integrated DNA Technologies (IDT). The forward primer used for library preparation was conjugated with 5’MMAE (IDT format: /5mod/; /5mod: NHS ester-PEG4-Val-Cit-PAB-MMAE (BroadPharm #: BP-25503 or ThermoFisher #NC3792832). Three reverse primers were synthesized with 5’-fluorescein. Although the primer sequences were the same, two primers (LJM-7540 and LJM-7542) were supplemented with 30 nt or 36 nt, respectively, 3’oligo-dA sequences separated from the priming sequence by an internal 9-atom triethylene glycol spacer (IDT format: /iSp9/). This spacer acts as a non-extendable linker facilitating separation of the aptamer strand from its complement during denaturing electrophoretic gel purification. (Supplemental Table S1). These three reverse primers were toggled throughout the selection to prevent inadvertent mutagenesis during selection rounds (Supplemental Figure S5).

The library template strand was synthesized with 5’ and 3’ terminal constant 20-nt primer regions flanking 40 random nt. All oligonucleotides were reconstituted at 100 μM. Primers were diluted to 5 μM. To create the 5’MMAE-conjugated naïve library for Round 1, PCR involved the following reagent volumes in a master mix: 640 μL 10× *Taq* polymerase buffer, 640 μL 1mg/ml (10X) BSA, 512 μL 50 mM MgCl_2_, 512 μL 2.5 mM dNTPs (dCTP, dATP, dGTP, and dTTP), 32 μL 100 μM library template DNA, 640 μL 5 μM forward primer, 640 μL 5 μM reverse primer, 2694 μL water, and 90 μL *Taq* DNA polymerase (Invitrogen # 10342020). Reagents were mixed, aliquoted into 100 μL volumes in PCR tubes, and subjected to thermal cycling using the following protocol: 95°C, 60 s; 6× (94°C, 15 s; 51°C, 35 s; and 72°C, 30 s); 4°C. Molar equivalent amounts of template and primers were used preserve library diversity and PCR cycles were limited to the minimum required to consume all 5’MMAE-conjugated forward primer.

PCR reactions were pooled and precipitated by addition of 2.5 volumes ethanol after addition of glycogen carrier and sodium acetate to a final concentration of 0.3 M. After washing with 70% ethanol and air drying, PCR products were resuspended in water and deionized formamide (Ambion #AM9342), heated at 90°C for five min, and subjected to electrophoresis through a 7.5% polyacrylamide gel [7.5 M urea, 19:1 acrylamide : bisacrylamide (Bio-Rad #1610144)]. As the forward primer was conjugated to MMAE (Supplemental Table S1), the forward strand migrated near the 90-nt marker, permitting purification of the aptamer library strand from its fluorescein labeled complement (Round 1, Supplemental Figure S5). Aptamer libraries were visualized by UV shadowing, excised with a razor blade, diced, and incubated in 0.3M NaOAc buffer at 56°C with agitation overnight. Eluted DNA was purified from gel pieces by passing through glass wool and ethanol precipitated as described above. The concentration of the resulting naïve library was determined with a Nanodrop instrument at 260 nm using a library-specific estimated extinction coefficient (757, 575 L/mol-cm). A 1 μM aptamer dosing solution in PBS containing 1 mM MgCl_2_ (280 pmol aptamer/280 μL) was assembled from the purified material.

### PDX Models

Short-term explants of PDX cells were obtained from the Mayo Clinic Brain Tumor Patient-Derived Xenograft National Resource. PDXs were then established directly from patient biopsy or surgical samples as flank tumors in athymic nude mice and maintained as previously described (20). These tumors were then used to establish short-term explant cultures or intracranial xenografts (20). Each PDX model is highly annotated with multi-omic characterization and relevant clinical data as previously described (21). Identities of PDX lines were verified by short tandem repeat analysis for each animal study.

### Mice

Female athymic nude (Strain #553) mice were purchased from Charles River Laboratories. All animal experiments were approved by Mayo Clinic Institutional Animal Car and Use Committee (IACUC), Protocol #A00006585-22. Orthotopic tumor inoculation and bioluminescence imaging were performed as previously described (6, 20).

### Cell culture

GBM PDX lines were maintained as previously described (21). Short-term explant cultures of GBM39, G39-eGFP/FLUC2, and G43 were grown in DMEM (Fisher Scientific #MT10013CV), supplemented with 10% fetal bovine serum (Gibco #220346) and 1% penicillin/streptomycin (Life Technologies #221674) at 37°C, 5% CO_2_. PDX lines were tested every 6 months for mycoplasma using a MycoAlert mycoplasma detection kit (Lonza #LT07-418). SVG-A was grown in DMEM (Fisher Scientific #MT10013CV), supplemented with 10% fetal bovine serum (Gibco #220346) and 1% penicillin/streptomycin (Life Technologies #221674) at 37°C, 5% CO_2_. HLF and NIH/3T3 (the gifts of A. Haak) were grown in DMEM/F12 (Gibco #11330-032) supplemented with 10% fetal bovine serum (Gibco #220346) and 1% penicillin/streptomycin (Life Technologies #221674) at 37°C, 5% CO_2_.

### *In vivo* SELEX in orthotopic PDX mice

Prior to SELEX, two cohorts of mice were injected with explants of GBM39-eGFP/FLUC2 cells (Supplemental Table S4). In the first cohort, 12 mice were injected in three groups (four mice per group) with the following cell counts: 100, 000 cells, 10, 000 cells, 1, 000 cells. In the second cohort, 12 mice were injected in four groups (three mice per group) with the following counts: 100, 000 cells, 50, 000 cells, 10, 000 cells, 1, 000 cells This was done to distribute tumor development over time. Bioluminescent Imaging (BLI) was performed to monitor tumor size over time and once readings ranging from 5×10^8^ to 1×10^9^ were observed, mice were injected with MMAE-conjugated aptamer library for each round of SELEX.

On the day of selection, one mouse received an intraperitoneal (i.p.) injection of 250 pmol aptamer library in 250 μL PBS containing 1 mM MgCl_2_. 4 h post-injection, the mouse was euthanized with CO_2_ and subjected to scrupulous transcardial perfusion with 30-60 mL PBS as follows: the euthanized mouse was pinned outstretched at a vertical angle and a 21g × 1” needle was inserted into the left ventricle while the heart was still beating. The mouse was perfused until the liver changed from dark red to pale brown with the effluent gravitationally pooling below the mouse. At each round, the brain was extracted, and the GFP^+^ tumor was dissected with the aid of goggles illuminating the specimen with UV light. Additional organs (adjacent brain, cerebellum, heart, lungs, liver, kidneys, spinal cord, and sciatic nerve) were also excised for the final three selection rounds. Bone marrow was also collected by centrifugation of both femurs at ≥10, 000 × g for the final three selection rounds. All tissues were snap-frozen on dry ice and stored at −80° C.

An optimized aptamer extraction method maximized yield relative to previous work in our lab (19). Bone marrow samples were rinsed with 500 μL PBS and subjected to centrifugation at 200 × g for 5 min to remove lysed blood cells and then processed as for the remaining tissue samples. Tubes with frozen tissue samples (≤ 200 mg) were thawed on wet ice and homogenized in 300 μL Qiagen Plasmid Extraction Kit Buffer P1 with a plastic pestle. Lysis buffer (300 µL: 50 mM Tris HCl, pH 7.4, 150 mM NaCl, 1% NP-40, 0.5% Na Deoxycholate, 1 mM EGTA, and 1 mM NaF) was then added and incubated on wet ice for 3 h. Samples were then sonicated at power 8 (Sonic Dismembrator 60, Fisher Scientific) for 10 s three times with 30-s of cooling on ice between sonications. Samples were then heated to 95° C for 10 min and subjected to centrifugation at 13, 100 × g for 5 min. The supernatant was removed and analyzed by qPCR as described above.

A 320-µL sample of this crude extract containing tumor-isolated aptamer was added to a 3.2 mL PCR reaction mix composed of 320 μL 10× Taq polymerase buffer, 320 μL 10× BSA, 256 μL 50 mM MgCl_2_, 256 μL 2.5 mM dNTPs (2.5 mM dCTP, dATP, dGTP, and dTTP), 320 μL 5 μM 5’MMAE-conjugated forward primer, 320 μL 5 μM reverse primer, 1043 μL water, and 45 μL *Taq* DNA polymerase (Invitrogen #10342020). From this and a separate 100 µL water control reaction, 5 aliquots (20 μL each) were used for cycle optimization. These 20-µL samples were removed at cycles 0, 25, 20, and 35 and the optimal number of cycles was determined by the homogeneity and yield of the PCR product band and consumption of primers analyzed after electrophoresis through a denaturing 10% polyacrylamide gel. The remaining volume was amplified by PCR for the optimal number of cycles determined for that round (typically ∼30 cycles). PCR reactions were pooled, precipitated from ethanol after addition of NaOAc and glycogen as before, and resuspended in water and deionized formamide. Samples were heated at 90°C for 5 min prior to loading on eight lanes a pre-run 7.5% denaturing polyacrylamide gel and subjected to electrophoresis at 600 V/22.86cm for 2.5 h. The forward MMAE-conjugated-labeled strand was distinguished from the FAM-labeled reverse strand by shadowing with a UV lamp and excised with a fresh razor blade (Supplemental Figure S5). This band was distributed between eight microcentrifuge tubes, finely diced, the pieces resuspended in 0.3M NaOAc buffer, and incubated overnight at 56°C with agitation. The eluant was passed through a 1.5 mL Eppendorf tube pierced with a hot needle and packed with glass wool to remove large polyacrylamide fragments. ApDCs were precipitated from the resulting filtered solution using ethanol as before and resuspended in 80-100 μL water. The absorbance at 260 nm (Nanodrop) was divided by the molar extinction coefficient to calculate the concentration of the recovered library. The sample for the next round of selection was prepared by adding 28 μL 10× PBS, 2.8 μL 100 mM MgCl_2_, 280 pmol of library, and increasing the volume to 280 μL with water. Samples were snap cooled by heating to 90°C for 5 min and cooling on wet ice for 10 min. Note that excess volume was prepared every round to ensure the mouse received an injection volume of 250 µL. This cycle was repeated for each round of selection.

### Aptamer sequencing

At Round 10, the library was initially sequenced with Azenta’s Amplicon-EZ sequencing to check for enrichment. The files were merged with the usearch - fastq_mergepairs function. The merge files were filtered using the usearch maxee function, discarding files with errors >0.5. Reverse strands were identified and the reverse complements created using SeqKit reverse complement tool. The two files were combined with the simple concatenate function in Linux. FASTAptameR2.0 was used to convert the resulting .fastq file for initially screening and evaluation of enrichment (22).

The naïve oligonucleotide library containing 5’MMAE-conjugated ApDCs, the libraries from 10 selection rounds, and libraries extracted from all tissues in the Round 10 mouse were subjected to deep sequencing. Gel-purified material was used in each case, including aptamers from Round 10 organs. Libraries were amplified for the number of cycles determined by analytical PCR in each selection round using unmodified primers. Duplexes were created from the libraries with PCR reactions containing: 10 μL 10× *Taq* DNA polymerase buffer, 10 μL 10× 1 mg/mL BSA, 8 μL 50 mm MgCl_2_, 8 μL 2.5 mm dNTPs, 10 μL library, 10 μL 5 μM forward primer, 10 μL 5 μM reverse primer, 32.6 μL water, and 1.4 μL *Taq* DNA polymerase. PCR reactions were thermal cycled for 6 rounds and then 20 µL fresh master mix was added to reach round and run for one cycle to ensure fully-extended duplexes (23). PCR product size and quality were confirmed by electrophoresis through an 8% native polyacrylamide gel (29:1 acrylamide:bisacrylamide; Bio-Rad #1610146).

PCR products were purified with a Qiagen MinElute Purification Kit (Qiagen #28004). Ten ng of eluted DNA, as quantified by the Qubit HS Duplex DNA Quantification Kit (Invitrogen #Q32851), were used as input for the NEBNext Ultra II DNA Library Prep Kit (NEB #E7645S) using associated NEBNext Multiplex primers index (primers set 1 and set 2; NEB, E7335S and E7500S). After preparing samples according to the manufacturer’s instructions (recommended volumes were halved and in the purification of adaptor-ligated DNA the Qiagen MinElute Purification kit was used instead of beads), samples were assessed by gel electrophoresis through an 8% native polyacrylamide gel (29:1 acrylamide:bisacrylamide, Bio-Rad #1610146) prior to pooling and analysis by high-throughput paired-end 150 cycle sequencing on a single lane of an Illumina MiSeq instrument (Mayo Clinic Sequencing Core).

Sequencing files were received as .cram files, which were then converted to .bam and subsequently .fastq files. Usearch was used to first merge paired-end reads. The merge files were filtered using the usearch maxee function, discarding files with errors >0.5. Reverse strands were isolated, and the reverse complements created using SeqKit reverse complement tool. The two files were combined with the simple concatenate function in Linux. AptaSUITE was used to filter any reads that did not contain both forward and reverse primers within an error of 3 bases, and to rank aptamers by sequence or cluster abundance (24).

Following deep sequencing analysis, aptamer enrichment was analyzed by identifying the most prevalent sequences and sequences clusters at Round 10 and by determining area under the curve of all sequences identified at Round 10. Biodistribution of individual ApDCs was estimated by:

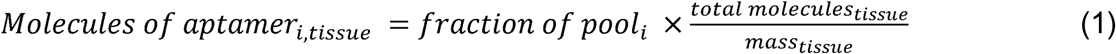

where *i* indicates a given individual aptamer, and *fraction of the pool_i_* is calculated by number of counts for aptamer *i* divided by the total of number of counts in the pool. The total number of molecules was determined by qPCR analysis of the library in every tissue sample. Recovered fraction of injected dose for aptamers was calculated by:

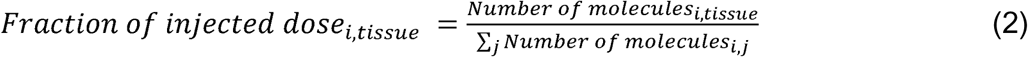

### Confirmation of library conjugation

To confirm prepared libraries were successfully conjugated to MMAE, 1pmol of ApDC library was incubated with or without anti-MMAE antibody (Creative Diagnostics # DMABA-JX124) at 1:10 dilution for 1 h at room temperature. Solutions were subjected to electrophoresis through 8% native polyacrylamide gels (29:1 acrylamide:bisacrylamide Bio-Rad #1610146) and stained with SYBR gold (Invitrogen #S11494).

### Synthesis of individual candidate ApDCs

Individual candidate aptamers were synthesized by IDT. ApDCs were created via PCR, scaling up the basic 100 µL reaction as needed to 1-4 mL: 10 μL 10× *Taq* DNA polymerase buffer, 10 μL 10× 1 mg/mL BSA, 8 μL 50 mm MgCl_2_, 8 μL 2.5 mm dNTPs, 10 μL library, 10 μL 5 μM 5’MMAE forward primer, 10 μL 5 μM 5’phosphorylated reverse primer, 32.6 μL water, 1 µL of 100 µM template and 1.4 μL *Taq* DNA polymerase. Following thermocycling, the PCR reaction was pooled, precipitated from ethanol and resuspended in 88 µL water plus 10 µL of 10× phage lambda exonuclease digest buffer and 2 µL of phage lambda exonuclease (New England Biolabs). The solution was incubated at 37°C for 30 min to hydrolyze the reverse strand. An equal volume of formamide was added to the reaction and followed by heating at 90°C for 5 min. ApDCs were subjected to electrophoresis through a 7.5% denaturing polyacrylamide gel at 600 V/22.86 cm for 2.5 h and were visualized by UV shadowing and excised from the gel with a razor blade. The gel fragment was minced and subjected to elution and ApDC purification proceeded as described above.

### Ex vivo tissue staining

Untreated mice bearing G39 tumors were euthanized and perfused with paraformaldehyde (PFA), followed by sucrose. Brains were excised and frozen on dry ice in optimal cutting temperature (OCT) compound. Tissues were cut into 5 μm sections. On the day of staining, OCT slides were dried for 10 min at room temperature. Sections were washed once with PBS for 5 min. Antigen retrieval was performed by incubating slides in boiling 10 mM citrate buffer, pH 6.0 for 30 min followed by washing with distilled water and PBS for 5 min each. A Pap Pen was used to trace the tumor. The tissue was pre-blocked in PBS containing 1 mM MgCl_2_, 2% BSA in PBS and 0.1 mg/mL sheared salmon sperm DNA for 1 h. The tissue was washed with binding buffer (PBS containing 1 mM MgCl_2_) twice for 5 min each. ApDC solutions (50 nM in PBS containing 1 mM MgCl_2_) were added to the tissue and incubated for 30 min. Slides were washed with binding buffer once followed by washing twice with PBS for 5 min each. Sections were then fixed with fresh 4% PFA for 15 min. Slides were blocked with 2% goat serum, 2% BSA in PBS for 1 h followed by an incubation with anti-MMAE antibody solution (Creative Diagnostics # DMABA-JX124; 1:500 dilution in 2% goat serum with 2% BSA in PBS) at 4°C in a humidified box overnight. Sections were washed with PBS 4 times. Anti-rabbit antibody conjugated to AF647 (Invitrogen #A32733) was then added at a dilution of 1:500 in 2% serum with 2% BSA in PBS for 1 h in the dark. Sections were washed 4 times with PBS and slides were then stained with 300 nM DAPI solution (Biolegend #422801) prepared in PBS for 5 m. After washing twice with PBS, slides were mounted using mounting media (Prolong glass antifade, Invitrogen #P36980). All steps were performed at room temperature unless otherwise stated. Slides were imaged on an LSM 980 microscope.

### ApDC binding to target cells by microscopy

Explants of G39-eGFP cells were seeded at 1×10^6^ cells per microscopy dish in (Mattek, #P35GC-1.5-14-C) overnight. Cells were ∼95% confluent on the day of the experiment. Cells were washed with wash buffer (PBS containing 1 mM MgCl_2_ and 4.5 g/L glucose) prior to incubation with 200 nM of heated (90°C 5 min) and snap-cooled ApDC in selection buffer (1 mM MgCl_2_ in PBS) with 4.5g/L glucose, 100 μg/mL sheared salmon sperm (Invitrogen #15632011V) and BSA (New England BioLabs #B9200S) for 30 m at 37°C. Cells were washed with wash buffer three times prior to fixation with 3.7% formaldehyde for 15 min. Cells were stained with anti-MMAE antibody (Creative Diagnostics # DMABA-JX124) 1:500 dilution for 1 h at room temperature followed by 3 washes with wash buffer and staining with secondary anti-rabbit antibody conjugated to AF647 (Invitrogen #A32733) diluted 1:500 for 1 h at room temperature. The cells were washed 3 times and stained with DAPI (Sigma-Aldrich, #10236276001). Cells were imaged with an LSM 980 confocal microscope.

### ApDC binding to cells as determined by qPCR

For cell binding determination by qPCR, SVG-A, HLF, 3T3, and explants of G39-eGFP and parental G39, cells were plated in 24-well plates at 60, 000 cells per well. Cells were washed with wash buffer (PBS containing 1 mM MgCl_2_) once. ApDC or aptamer solution at 50 nM in PBS containing 1mM MgCl_2_ was heated at 90°C for 5 min and cooled on ice for 10 min. BSA and sheared salmon sperm DNA were added, each to a final concentration of 0.1 mg/mL, as well as glucose to a final concentration of 4.5 g/L. The ApDC or aptamer solution was added to the cells and incubated at 37°C for 30 m. Following binding, the cells were washed at room temperature three times with wash buffer. Cells were recovered by scraping on ice in 100 µL wash buffer. Wells were washed once with a second portion of 100 µL wash buffer to collect any remaining cells. Collected material was subjected to centrifugation at 500 × g for 5 m and the supernant aspirated. Equal volumes of lysis buffer (50 mM Tris HCl, pH 7.4, 150 mM NaCl, 1% NP-40, 0.5% Na Deoxycholate, 1 mM EGTA, and 1 mM NaF) and Qiagen Plasmid Extraction Kit Buffer P1 were added to the cells and vortexed to resuspend. The lysate incubated on ice for 30 m, prior to sonicating each sample for 10 s at power 8. Lysates were heated at 95°C for 10 min and then subjected to centrifugation at 13, 100 × g in a microcentrifuge. The supernatant was used for qPCR (QuantaBio PerfeCTa #95071-012). Protein was quantified as determined by Qubit Protein Assay Kit (Invitrogen #Q33211).

### Biodistribution of individual aptamers

Mice were stereotactically injected with 50, 000 explanted GBM39-eGFP cells, as described above. Mice were closely monitored for weight and behavioral changes as signs of tumor progression. Animal approaching moribund status were enrolled in the study. ApDC cocktail solutions of 350 pmol/350 μL (50 pmol each ApDC or unconjugated aptamer) were created in PBS containing 1 mM MgCl_2_, heated for 5 min at 90°C and rapidly cooled on ice for 10 min. The random region for Aptamer 7 is identical to that of negative control X from Doherty et al. 2025 (19). Control vehicle solutions lacking aptamers were created at the same time. Mice were injected i.p. with 350 μL aptamer cocktail (350 pmol) or vehicle. Four h post injection, mice were euthanized and underwent scrupulous transcardial perfusion as described above. For biodistribution experiments a right intracardiac perfusion was done followed nicking of the infrarenal abdominal aorta to ensure blood removal from the liver and kidneys. The brain, heart, lungs, spleen, liver, kidneys, bone marrow, and sciatic nerve were collected. The GFP^+^ tumor was carefully excised with visualization by UV illumination. The bone marrow was again collected from both femurs by centrifugation at ≥10, 000 × g. All tissues were rapidly frozen on dry ice and stored at −80° C.

Aptamers were extracted from tissues as described above. qPCR was performed using the library forward primer and sequence-specific reverse primers to quantitate individual aptamers (Supplemental Tables S1-S3).

### ApDC pharmacokinetics

For pharmacokinetic analysis, ApDC cocktail solutions contained two selected ApDCs and the negative control ApDC. Mice were injected with 50, 000 G39-eGFP explant cells. Mice were closely monitored by BLI, weight, and outward behavior to judge tumor growth. On the day of the experiment, mice received an I.V. tail vein injection with 100 µL ApDC cocktail solution (1.5 µM, 50 pmol per individual ApDC). Mice were euthanized at either 0.5 h, 2 h, 4 h, 6 h, or 8 h, thoroughly perfused through the right ventricular intracardiac route, followed by clipping the infrarenal abdominal aorta and continued perfusion. Tissues were processed and qPCR was performed as described above.

For ApDC extraction from serum, lysis buffer (170 µL: 50 mM Tris HCl, pH 7.4, 150 mM NaCl, 1% NP-40, 0.5% Na Deoxycholate, 1 mM EGTA, and 1 mM NaF) was added to 30 µL of mouse serum on ice and vortexed. Samples were heated to 95°C for 25 min and the denatured protein was pelleted by centrifugation at 13, 100 × g for 5 min. The supernatant was removed and further diluted with water (300 µL) before the sample was subjected to phenol: chloroform extraction twice to remove remaining protein. The aqueous extract was ethanol precipitated as described above and the sample was resuspended in 30 µL of water and used for the qPCR reaction as described above.

## Results

### Evaluating 5’MMAE-conjugated primers for SELEX

Striving to mimic ADC technology by substituting the antibody targeting moiety with an unmodified DNA aptamer, we utilized the common ADC linker and toxin construct of MMAE-PAB-VC (Figure 1A). Trans-cyclooctene (TCO) – tetrazine (Tz) click chemistry was initially tested for conjugation of MMAE-PAB-VC-PEG4-Tz to a 5’TCO DNA primer, which was confirmed by gel mobility retardation upon electrophoresis through a 16% polyacrylamide denaturing gel (Figure 1B). A challenge was batch-to-batch variation in the reactivity of synthetic 5’TCO DNA oligonucleotides, hypothesized to be due to varying degrees of trans-to-cis isomerization of the cyclooctene group, the latter being unreactive (Supplemental Figure S2). After inconsistent yields in multiple pilot experiments, commercially available 5’MMAE DNA primers synthesized by conventional 5’amino-DNA oligonucleotide reaction with an MMAE-N-hydroxysuccinimide (NHS) derivative were used for DNA conjugate library preparation for convenience.

In our proposed *in vivo* selection strategy, aptamer ‘fitness’ is defined by anti-GBM aptamers that can carry cargo to the tumor, and the “reward” is PCR amplification. We hypothesized, based on previous work (Supplemental Figure S1), that toxin-bearing aptamers should be selected from a toxin-bearing random library. Thus, the ApDC library must initially carry MMAE and be restored with MMAE conjugation at each selection round. To ensure that ≥95% of the library was conjugated with 5’MMAE, we elected to use a 5’MMAE conjugated primer in library PCR followed by gel purification. We therefore confirmed that MMAE is stable to PCR conditions. Free MMAE was subjected to 20 cycles of PCR thermal cycling and was then compared with untreated MMAE using LC-MS/MS as previously described in Jain et al. (Figure 1C) (25). No mass difference was detected between the control and PCR-treated MMAE samples, indicating the MMAE was structurally stable through repeated rounds of heating to 95°C and cooling to 50°C. To further support this, we tested the biological activity of free MMAE following 25 cycles of PCR thermocycling followed by heating at 95°C for 5 min and rapid cooling to simulate PCR conditions. No loss of MMAE toxicity was observed after thermocycling, further supporting the thermal stability of MMAE (Supplemental Figure 3B).

To confirm that Taq DNA polymerase could incorporate a 5’MMAE-modified DNA primer during conventional PCR, two reactions were assembled based on SYBR green detection: one with a 5’MMAE-modified primer and the other with a standard primer. Aliquots were removed at increasing PCR cycles these samples were analyzed by native polyacrylamide gel electrophoresis. SYBR green fluorescent signal detects duplex DNA amplicons. Yields were equal in both reactions, accompanied by a detectable gel mobility retardation in reactions containing the MMAE-conjugated primer (Supplemental Figure 3B). Subsequent gel staining SYBR gold to evaluate both single strands and duplexes revealed both the anticipated gel mobility retardation of MMAE-conjugated product amplicons as well as the disappearance of the 5’MMAE-conjugated forward primer, confirming its successful incorporation into the PCR product by Taq DNA polymerase (Figure 1D).

### In vivo selection of MMAE-conjugated ApDCs in an orthotopic PDX GBM mouse model

Based on our previous *in vivo* SELEX experience with unconjugated DNA (19), we used a similar strategy but substituted an MMAE-conjugated DNA ApDC library, where MMAE served as a generic example cargo. *In vivo* SELEX was performed as outlined in Figure 2A. Athymic nude mice were implanted with explants of GFP/luciferase-tagged PDX line GBM39-eGFP/fLuc2, which originated from a 51-year-old male with a confirmed left frontal IDH-wild type GBM tumor as described (20). Mice were inoculated with varying numbers of tumor cells to stagger tumor growth enabling sequential selection rounds over time. GBM growth was followed by BLI (Supplemental Table 2). Once a BLI reading of 5×10^8^-1×10^9^ log total flux units was observed, a mouse was eligible for the *in vivo* selection protocol. On the day of selection, one mouse was injected intraperitoneal (i.p.) with 250 pmol ApDC library (∼10^14^ aptamers). ApDCs were allowed to circulate for four hours with the goal that rare, folded aptamers would reach the brain parenchyma, cross the BBB, and accumulate in the GBM tumor by unspecified mechanisms. The mouse was then euthanized and thoroughly perfused via left ventricular transcardial perfusion with saline to scrupulously remove aptamers present in circulation. The GBM tumor was then isolated guided by GFP signal, and GBM-associated aptamers were isolated by an optimized extraction method as previously described (19). Selection rounds were aborted if, at the time of harvest, the mouse was found to have a significant GBM hemorrhage. Recovered ApDCs were amplified by PCR again incorporating 5’MMAE conjugated primers. As a precaution against unexpected evolution of library size by deletion or insertion mutagenesis, we toggled the length of the unextendible 3’ tail of the reverse primer among three sizes, enabling clear identification of the desired library species by denaturing gel electrophoresis (Supplemental Figures S4-S5). Ten successful rounds of selection were completed. As a precaution, successful MMAE library conjugation was confirmed in the initial and final libraries (Figure 2B), and the apparent length of the conjugated ApDC library and derived duplexes submitted for sequencing were confirmed by electrophoretic analysis (Supplemental Figures S6A-B).

**Figure 2.**
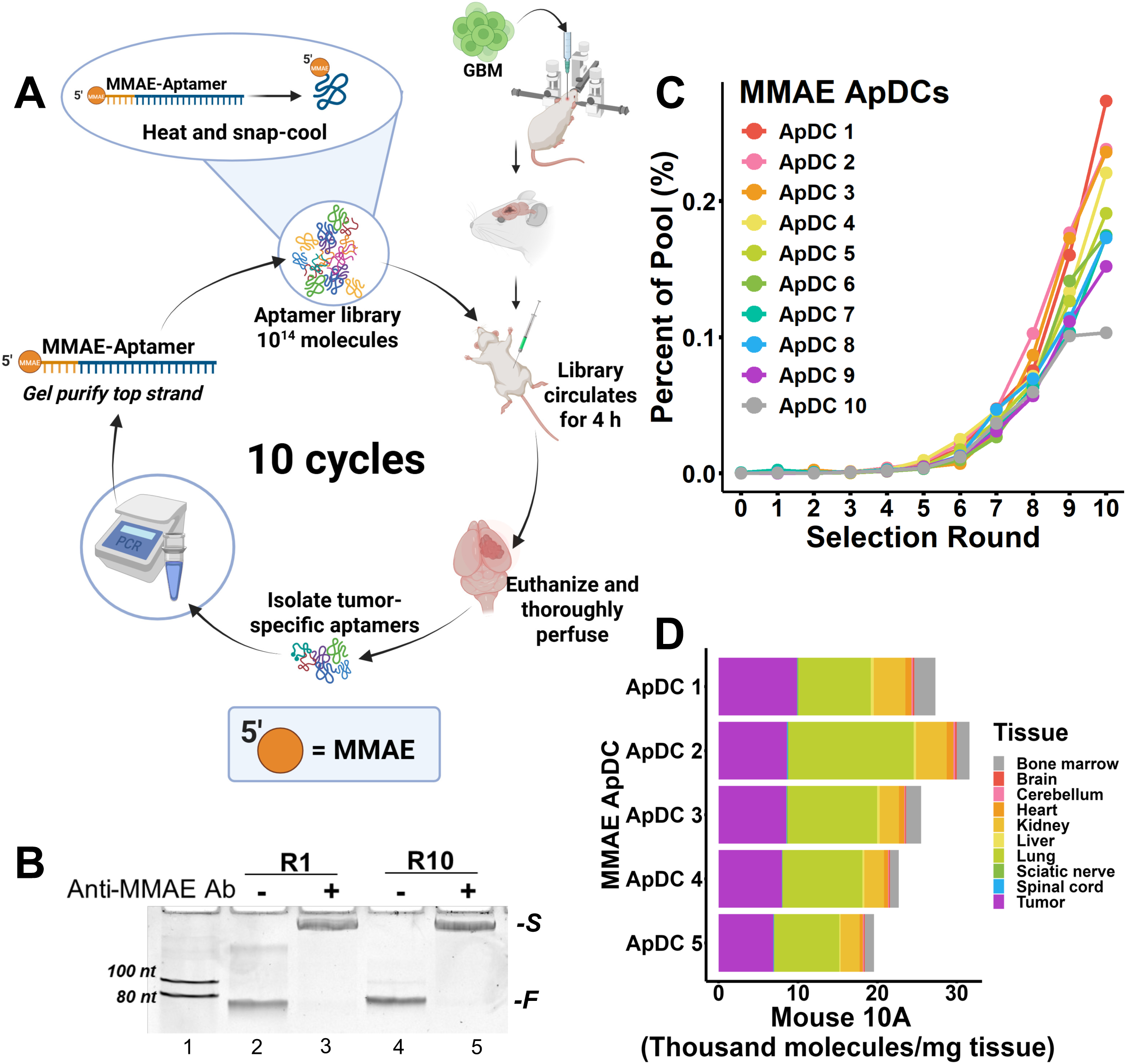
*In vivo* selection of ApDCs in an orthotopic patient derived xenograft (PDX) model. A. A schematic diagram of the *in vivo* SELEX with an ApDC library. B. Native polyacrylamide gel shift assay detecting the presence of MMAE with an anti-MMAE antibody in the naïve library used for Round 1 and the final library used for Round 10. *-F* = free library, *-S* = shifted library due to anti-MMAE antibody-ApDC complex C. Top 10 sequences by area under the curve at Round 10 from NGS. D. Biodistribution of the top 5 sequences identified at Round 10 (Figure 2C) in the Round 10 mouse.

After the final SELEX round, samples of adjacent normal brain, cerebellum, heart, lung, liver, kidney, bone marrow, spinal cord, and sciatic nerve were collected from the Round 10 mouse. GBM-homing libraries from all selection rounds and aptamers collected from all organ specimens from the Round 10 mouse were analyzed by bar coding and next generation sequencing. Selection progress could be monitored approximately by qPCR analysis of aptamers recovered from each tissue, but, as in our previous *in vivo* SELEX experiments, this analysis was not sufficiently informative to support detailed conclusions (Supplemental Figure. S7) (19). After deep sequencing and bar code deconvolution, aptamer sequences were sorted by GBM abundance at Round 10 and their percentage in the total sequenced GBM pool was graphed over the 10 selection rounds (Figure 2C). Exponential enrichment was observed among the leading sequences over the 10 SELEX rounds, indicating that selection was occurring within the library.

Analyzing the sequencing data further, the most abundant sequences were evaluated for their biodistribution in the Round 10 mouse as a measure of GBM homing specificity (Figure 2D). For many of the leading ApDC candidates, the GBM tumor was indeed a predominant tissue for aptamer homing. Interestingly, when compared with aptamer biodistributions of a library resulting from *in vivo* SELEX in the same model using an unconjugated DNA library (19), there was an obvious increase in the recovery of ApDC molecules from non-GBM tissues, especially the lung (Supplemental Figure S8A-C).

To further explore the increased amount of ApDC detected lung tissue, binding of the top five ApDC candidates (ApDC 1-5) was evaluated by qPCR in vitro vs. a negative control ApDC (scrambled version of ApDC 1, designated ApDC 6) using the target G39-eGFP cell line compared to normal human and mouse lung fibroblasts (Supplemental Figure S8D). We note that lung fibroblasts are simply a convenient surrogate for the wide variety of lung cell types. Selected anti-GBM ApDCs did not bind better to lung fibroblasts than to GBM cells *in vitro* (Supplemental Figure S8D). In fact, ApDC 2, ApDC 3, and ApDC 4 bound with higher affinity to the target G39-eGFP cells than to lung fibroblasts. It is possible that i) ApDC accumulation in lung may reflect greater physiological perfusion or reduced washout, or ii) it has been reported that free MMAE biodistribution favors well-vascularized tissues, with the lung having the highest area-under-the curve (AUC) and the largest tissue-to-plasma partition coefficients in PK analysis (26), This raises the possibility that unspecified receptors for the MMAE cargo contribute to ApDC biodistribution in addition to aptamer receptors.

We further evaluated the recovered Round 10 library to determine if this biodistribution pattern was a characteristic of the entire library. We identified a subset of ApDCs, many of which were in low abundance in the pool, that demonstrated greater accumulation in the GBM compared to any other tissue (Supplemental Figure S9A). Although these ApDCs were found to have favorable biodistributions, the leading example was ApDC 1, which, paradoxically, fails to detectably bind target GBM cells *in vitro* (Supplemental Figure S9B-D; Figure 3; Supplemental Figure S13-15). We hypothesize that ApDCs of this type selectively home to the tumor vicinity without direct GBM cell binding, perhaps recognizing tumor-specific elements of the extracellular matrix, tumor-recruited host stroma, or even antigens unique to endothelial cells of the GBM-induced vasculature.

**Figure 3.**
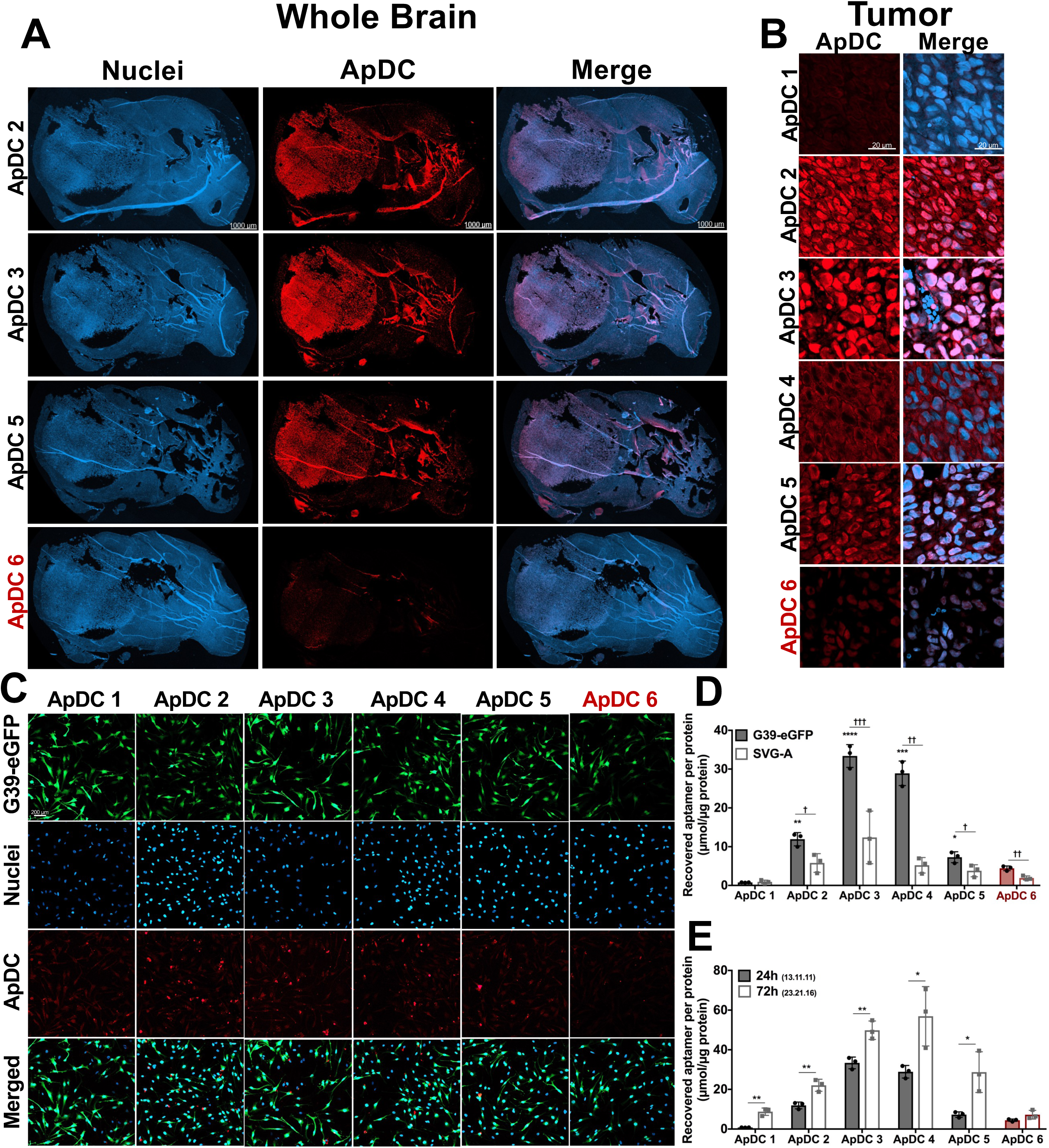
Evaluation of target tissue and cell binding. A. Ex vivo ApDC (50 nM) staining of serial sections flash frozen and OCT fixed from one untreated mouse bearing G39 orthotopic tumors (n= 2). B Higher magnification of ApDC staining in tumor in 3A. C-D. ApDC binding at 50 nM to target G39 cells in culture as detected by microscopy (C) and qPCR (D) Cells were incubated with ApDC 24 h after seeding. E. ApDC binding to G39 cells after 24h or 72 h after seeing the cells. Scale bars are 1 mm, 20 µm, and 200 µm respectively. Comparisons are to the negative control unless otherwise specified. *p<0.05; **p<0.01, ***p<0.001, and ****p<0.0001.

### In vivo-selected ApDCs bind GBM tumor tissue ex vivo and GBM cells in vitro

The top 5 ApDC candidates identified in the GBM tumor in SELEX Round 10 were selected for further study. We tested these conjugates for the ability to selectively bind GBM tumor tissue ex vivo in serial sections of untreated mouse brains bearing the G39 tumor. This histological approach, analogous to immunofluorescence, produced compelling positive results both for fresh frozen tissue and FFPE-preserved tissue (Supplemental Figure S10). All five ApDCs selectively stained GBM tissue relative to negative control ApDC 6 (Figure 3A, Supplemental Figure S11). ApDC 2, ApDC3, and ApDC5 stained the GBM tumor particularly brightly relative to normal brain, confirming tumor specificity. In contrast, ApDC 1, with a signal only slightly above the negative control ApDC, stained the GBM tumor poorly relative to the other candidates. ApDC 4 demonstrated strong but patchy staining within the tumor and appeared to bind normal brain issue more than other candidates. In one tissue study, the GBM tumor was removed during sectioning, prior to staining. Thus, only normal brain tissue was present. These tissue sections demonstrated no signal from any of the ApDC candidates. Staining was observed in further serial sections from the same mouse where GBM tumor was retained, further supporting GBM-specificity of the *in vivo* selected ApDCs (Supplemental Figures S12-S13).

*Ex vivo* GBM staining shown in Figure 3A was further evaluated at higher magnification (Figure 3B). Interestingly, ApDCs were characterized by different *ex vivo* staining patterns, supporting the hypothesis that they target different tumor antigens. ApDC 2 showed strong nuclear and cytoplasmic staining, whereas ApDC 3 staining was predominantly nuclear with some perinuclear staining. ApDC 4 staining was cytoplasmic. ApDC 5 staining was primarily perinuclear with some nuclear staining, but staining was less intense than for ApDC 3. Although weak signals had been observed at low magnification for ApDC 1, examination of ex vivo-stained GBM cells at higher magnification did not reveal any ApDC 1 staining, further supporting the possibility that ApDC1 targets host elements in tumor microenvironment not uniformly present in tissue sections. Thus, four of the five selected ApDC candidates bound GBM tissue ex vivo, suggesting additional applications for these reagents.

Because GBM tumors *in vivo* involve many cell types including host cells and host vasculature, we evaluated selected ApDC binding to the purest GBM cells available to us, G39 cells grown in culture. Similar to *ex vivo* tissue section staining results, ApDCs 2-5 demonstrated binding to target GBM cells by microscopy and qPCR, while ApDC 1 did not (Figure 3C-D). To extend the results obtained for *ex vivo* staining of mouse brain and tumor sections, we tested binding of the selected ApDCs to target GBM cells relative to a normal human astrocyte cell line, SVG-A (Figure 3D). ApDCs 2-5 all bound to the target GBM cells significantly more than the normal human astrocytes. ApDCs 2, 4 and 5 bound SVG-A comparably to the negative control ApDC, supporting the specificities of these selected ApDCs. Interestingly, ApDC binding to cultured target GBM cells detected by indirect immunofluorescence microscopy is weaker than *ex vivo* staining of tumor tissue sections using the same method. This suggests that the target ligands for these ApDCs are either i) upregulated *in situ*, ii) presented at higher density in intact tissue, and/or iii) secreted to accumulate in the intact tumor microenvironment.

It is known that GBM cells transplanted into mice display a wider variety of cell phenotypes than in culture. It is thus possible that the target ligands of these ApDCs are reduced in cell culture. It was also noted that ApDC binding signal increased with prolonged culture times and differed for GBM cells from different passages, supporting that the ApDC targets are variable and possibly inducible (Figure 3E).

In one experiment GBM cells cultured for 72 h post-seeding prior to ApDC binding analysis were treated with trypsin to evaluate if increased ApDC binding signal was attributable to extracellular matrix (ECM) deposition (Supplemental Figure S14A). Only ApDC 3 demonstrated significant reduction in cell binding upon trypsin treatment. This could indicate deposition of this target ligand in the ECM or ApDC recognition of a trypsin-susceptible protein target.

Finally, we sought to evaluate the ability *in vivo* selected ApDCs to tolerate cargo toxin substitution. We first simply removed MMAE from the selected ApDCs and monitored binding of the unconjugated aptamers to cultured GBM cells (Supplemental Figure S14B). Surprisingly, none of the aptamers tolerated this change when cultured cell binding was quantified by qPCR. We then tested the substitution of MMAE with its very similar sister toxin, MMAF, which differs from MMAE only by the presence of one terminal carboxyl group. Again, this substitution was not tolerated as judged by cultured GBM cell binding assays (Supplemental Figure S14C). As the *in vivo* selected candidates clearly bind the target cells and tissue above that of an untrained sequence, but binding to the target cells in culture is extremely sensitive to the presence and character of the conjugated cargo, we conclude that the selected ApDCs are highly sensitive to the cargo in their folding and/or target engagement. Secondary structures predicted by mFold (27) do not provide insight (Supplemental Figure S15) (19). The observed extreme sensitivity to post-selection conjugate modification was not previously observed in studies of cell- or protein-selected aptamers. We hypothesize that this characteristic is specific to *in vivo* SELEX. This result again emphasizes the conclusion that selection libraries carry the specific conjugate of interest, not assuming that aptamer targeting function will tolerate cargo changes post selection.

### Robust biodistribution prediction from deep sequencing data

We sought to evaluate independently the biodistribution of the top 5 ApDCs from the *in vivo* SELEX experiment, allowing comparison with deep sequencing data obtained from Round 10. To do so, we synthesized a cocktail of the 5 most prevalent ApDCs plus negative control ApDC 6, and an unconjugated negative control sequence, Ap 7 (Figure 4A). Mice were injected with 350 µL of cocktail (1 µM; 50 pmol per sequence) or vehicle, and the biodistribution was determined in collected tissues 4 h post-I.P. injection using qPCR with aptamer-specific primers. All 5 trained ApDC sequences were found to accumulate in the brain tumor significantly more than either the conjugated or unconjugated sequences, at levels of ≥ 10-20×10^6^ molecules/mg of tissue. This is significantly higher than vehicle background (Figure 4B). Negative controls ApDC 6 and Ap7 were detectable in the GBM tumor, an observation that we hypothesize to be due to BBB leakiness in the G39 model. Evaluating other tissues including the adjacent uninvolved brain and cerebellum indicated that the levels of aptamer accumulating in these tissues are exceedingly small relative to the GBM tumor: at most, some of the selected ApDCs accumulate ∼0.5×10^6^ aptamers/mg of tissue (Figure 4C). Interestingly, ApDCs 1 and 5 are not different by qPCR detection from negative control sequences found in adjacent normal brain and cerebellum, supporting the GBM-selectivity of these ApDCs.

**Figure 4.**
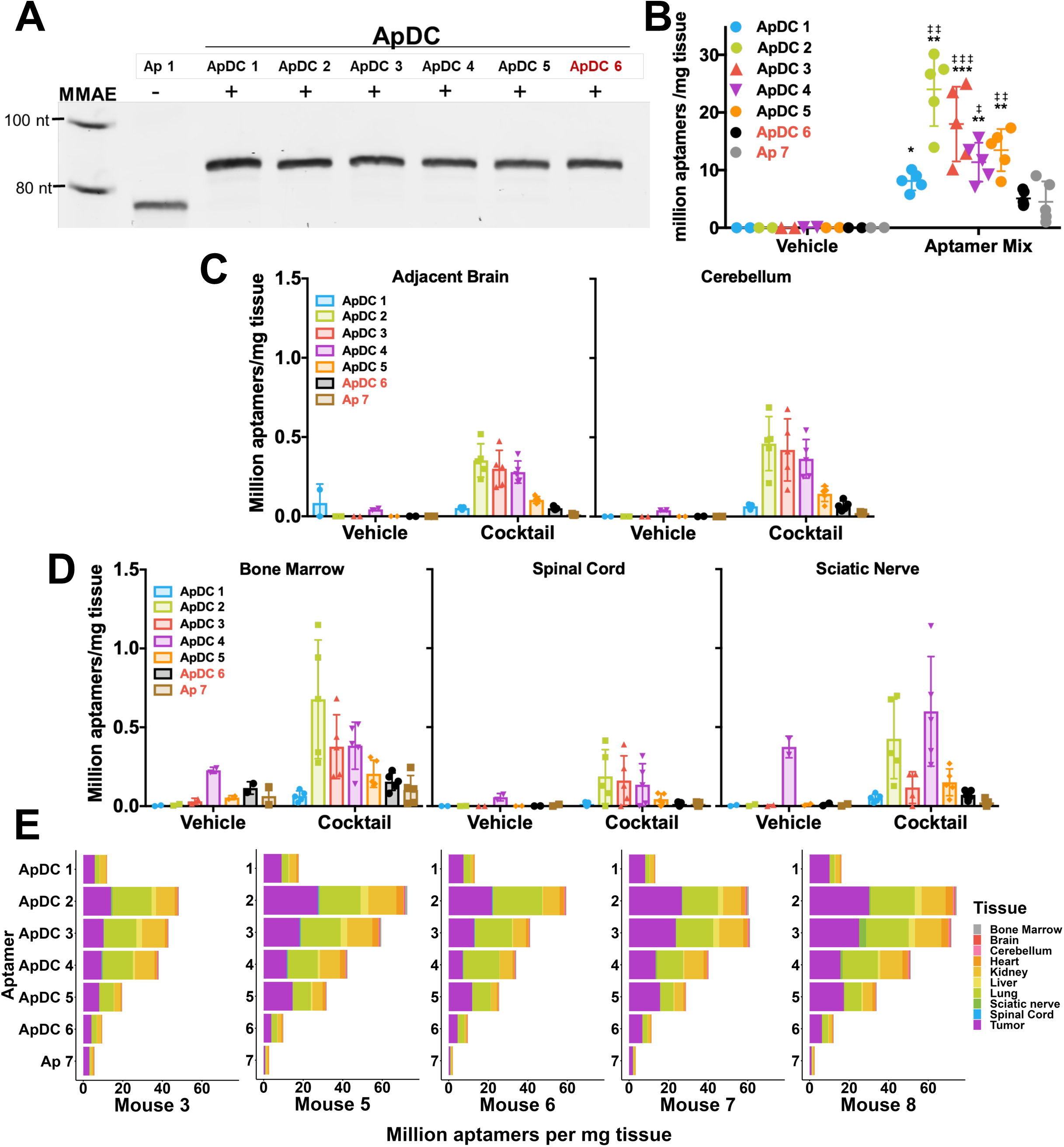
Biodistribution and *in vivo* specificity of the top 5 ApDC candidates. Mice were injected I.P. with 350 µl of 1 µM ApDC/aptamer cocktail (50 pmol per sequence) or vehicle. The cocktail was composed of the 5 top ApDC candidates, a scrambled sequence of ApDC 1, and the negative control aptamer sequence used in our previous *in vivo* SLELEX. 4 h post-injection, the mice were euthanized and thoroughly perfused prior to organ harvest. Tumor was detected by GFP positive staining. Aptamers were isolated from organs and detected with sequence specific PCR primers. A. Denaturing polyacrylamide gel stained with SYBR gold confirming the size shift and purity of the ApDCs in the cocktail injected into the mice. B. The amount of ApDCs and negative control aptamer detected in the GFP positive brain tumor. C. Comparison of the levels of ApDCs detected in the brain tumor compared to other areas of the brain. D. Levels of aptamer detected in tissues sensitive to anti-tubulin agents like MMAE. E. Whole organ biodistribution of the candidates across 5 mice. *= comparison to ApDC 6. ‡ = comparison to Ap 7. *p<0.05; **p<0.01, ***p<0.001, and ****p<0.0001.

We reviewed ApDC delivery to non-target tissues of particular concern for unwanted toxin delivery including bone marrow, spinal cord, and sciatic nerve (Figure 4D). Similar to normal brain, the ApDC signal in these tissues was exceedingly low, not accumulating more than 1×10^6^ molecules/mg tissue. Notably, some signal was detected in vehicle controls from these tissues, indicating inevitable low-level nonspecific background contamination during the greater handling and processing, especially for bone marrow. Finally, tissue biodistribution across all organs in all five mice was evaluated (Figure 4E). The tumor was highly enriched for all *in vivo* selected ApDCs. Signals for ApDC1 and 5 accounted for more of the remaining ApDC in the mouse at 4h than any other organ. As predicted from the deep sequencing results, lung and kidney did demonstrate significant ApDC accumulation at 4h.

To evaluate whether *in vivo* selected ApDCs might target other GBM or high-grade glioma models, we tested the ability of ApDC 2, 3, and 5 to home to either a different orthotopic mouse GBM (GL261) or a mouse diffuse midline glioma (DMG; MPP) comparing to negative control ApDC 6 (Supplemental Figure S16). Interestingly, all three *in vivo* selected ApDCs detectably accumulated in the brain and tumors more than negative control ApDC 6, but tumor specificity was not observed. We hypothesize that *in vivo* selected molecules retained their ability to home to the brain, accumulating in significantly higher amounts than in the orthotopic G39 GBM model, but that the specific G39 tumor ligands were absent in the alternative tested models so ApDCs lost tumor specificity. This result underscores the extreme specificities of the selected ApDCs.

### Pharmacokinetic study of ApDC 2 and 5

We wished to explore the pharmacokinetics of *in vivo-*selected ApDCs to better understand the accumulation and persistence of *in vivo-*trained ApDCs (Figure 5). We were also curious if ApDCs accumulated in the lung would wash out of this highly vascularized tissue over time. For this experiment, ApDCs 2 and 5 were selected for further study. ApDC 2 was selected as it accumulated more in the tumor *in vivo* than any other aptamer and also bound well to the GBM target cells in culture. ApDC 5 was selected because it bound both the target cells and tissue and demonstrated the least off-target tissue accumulation. The injection route was altered from i.p. to i.v. for comparison. We began by comparing the amount of ApDC accumulation in the GBM at 4 h (Figure 5A). Surprisingly, for ApDCs 2 and 5, injection route made no difference in ApDC accumulation in the orthotopic GBM 4 h post-injection. However, administration route showed an effect for negative control ApDC 6. We hypothesize that this is because *in vivo* selected aptamers were trained to perform well upon i.p. injection (Figure. 5A). Sustained ApDC 2 and 5 delivery (vs. negative control ApDC 6) is observed in all organs, especially in the orthotopic brain tumor (Figure 5B-C). By ∼ 4 h ApDC 2 and 5 concentrations plateau in the GBM, whereas negative control signal continues to decline, demonstrating the impact of *in vivo* SELEX incubation time on the properties of the selected molecules. The concentration of ApDCs in the lung remains high over 8 h. Interestingly, negative control ApDC 6 is slowly eliminated from the lung, a behavior not observed for the ApDC candidates, suggesting that lung accumulation may not be intrinsic to all MMAE-conjugated ApDCs, but reflecting properties of the conjugate and/or the mode of selected ApDC tissue delivery. In any case, the orthotopic GBM remains among the top organs accumulating *in vivo* selected ApDCs. Further, ApDC concentrations accumulated in the orthotopic GBM remain folds higher relative to normal brain tissue and sensitive organs (Figure 5C).

**Figure 5.**
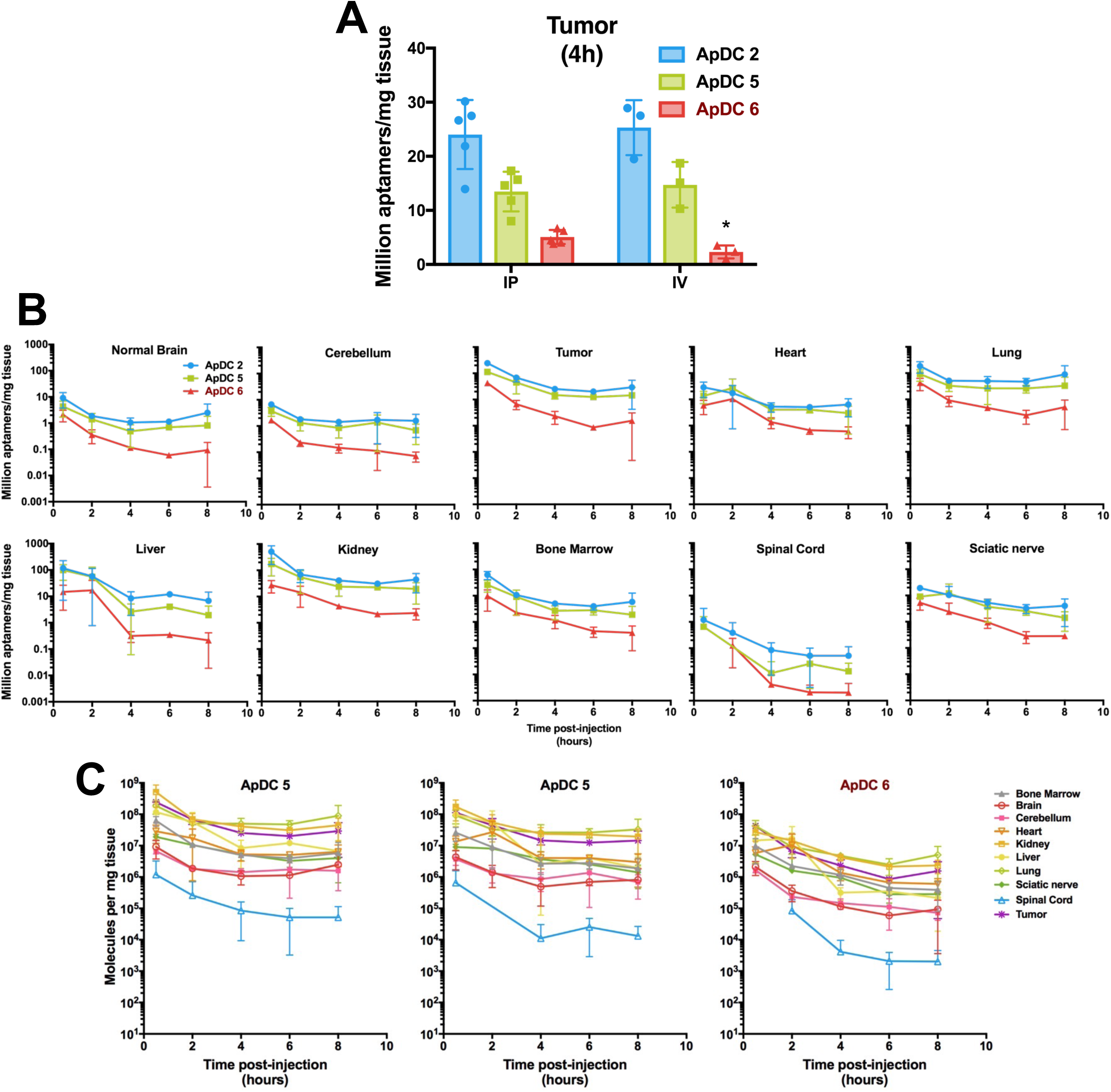
Pharmacokinetic study of *in vivo* selected ApDCs. Mice were injected I.V. with 100 µl of 1.5 µM ApDC cocktail (50 pmol per sequence) and harvested at either 30 min, 2h, 4h, 6h, or 8h post-injection. Mice were thoroughly perfused prior to organ harvest. Tumor was detected with GFP positive staining. Aptamers were isolated from tissues and detected with sequence specific PCR primers. A. Comparison of accumulation of ApDC in the tumor at 4h post- I.P. or I.V. injection. B-C. Detection of ApDC in serum (B) or other tissues (C) over 8h. *p<0.05; by two-sided *t*-test.

## Discussion

ADCs have demonstrated promise in high grade gliomas, such as GBM (5, 6, 8). However, their efficacy in GBM is in large part limited by the IgG antibody targeting moiety due to its relatively large size and inability to effectively cross the BBB, off-target toxicity, and limited tumor antigen targeting. Regardless of the current limitations of ADCs for GBM, there is increasing evidence that ADCs rarely achieve a higher maximum tolerated dose in humans than conventional therapies and only 0.1-2% of administered ADCs reach the tumor (28). Thus, the concept of these “biological missiles” is intriguing, but the selection of the antibody targeting moiety still rests entirely on rational selection of an antigen target. This points to the potential opportunities afforded by *in vivo* aptamer selection, where the targeting agent is smaller, may cross the BBB, and is developed by an evolutionary process for optimal *in vivo* performance without the need to specify a molecular target. The opportunity may even exist to treat with cocktails of anti-GBM aptamer to target multiple tumor antigens simultaneously.

Traditionally, unmodified aptamer libraries are subjected to *in vitro* or *in vivo* selection and then it is assumed or hoped that post-SELEX conjugation will not disturb aptamer specificity and affinity, such as the example provided by pegaptanib (29, 30). Post-SELEX modifications have tended to be small base modifications that increase the stability, pharmacokinetics, or pharmacodynamics of the molecule. Selections performed with a conjugated aptamer libraries are termed conjugate-SELEX and have been used to identify aptamers that function as successful targeting moieties (31). A major challenge with conjugate-SELEX, or any SELEX that incorporates substantial modifications, is that the modifications must be enzymatically and physically compatible with PCR (32–34). Given the failure to preserve cell specificity in our initial attempt to conjugate an ADC linker and toxin to DNA aptamers selected from unconjugated libraries *in vivo* (19), we hypothesized that conjugate-SELEX would be necessary to identify *in vivo-*relevant, anti-GBM ApDCs.

Our use of unmodified DNA undoubtedly limits the lifetime of the selected molecules. A unique outcome from this selection strategy is that ApDCs are quickly cleared from most tissues but accumulate in the tumor tissue remarkably well. The selected aptamers likely accumulated in the tumor as a result of the selection requirement of survival for 4 h post-injection. Due to the 4 h survival requirement, our approach rewarded long-lived ApDCs but missed molecules that targeted pathways which resulted in rapid intracellular or extracellular degradation. We note that both the SELEX process and our PCR-based analysis strategies detect only substantially intact aptamers.

In our prior study implementing *in vivo* SELEX with unconjugated DNA molecules (19) precautionary steps were taken to avoid triggering toll-like receptor 9 (TLR9) with unmethylated DNA 5’-CpG motifs (35). However, synthesis costs for the required aptamers were high and we did not observe significant benefit in these prior short-term experiments. Thus, the present *in vivo* SELEX library involved primer regions largely devoid of CpG motifs (there is one CpG in the reverse primer) and we hypothesized that *in vivo* SELEX would deplete from the pool sequences with problematic motifs in the random region. We note that two of the top candidates contain single CpG dinucleotides derived from the random region, but this motif is absent from other sequences. We further note that the cGAS-STING pathway is known to trigger innate immune responses to cytoplasmic DNA, but it is thought to be most sensitive to relatively long double-stranded DNA fragments (unlike the aptamers present in our libraries) (36). It remains possible that aptamers triggering the cGAS-STING pathway in tumor cells might be beneficial, as such effects might help to enhance an immune response. Future studies will be needed to address whether unmodified DNA aptamer libraries have the potential to activate the cGAS-STING pathway over extended time periods.

In planning our approach, we also selected a cargo conjugation strategy that would maintain library diversity while enabling nearly 100% library conjugation. By demonstrating the function of MMAE-conjugated forward primers during PCR, all recovered ApDC molecules should have the ability to compete and be rewarded at each round of selection. An alternative option that was considered was to conjugate MMAE to the library after PCR amplification. We were concerned that the efficiency of conjugation could be more difficult to determine and aptamer folding might impose conjugation bias at this step. Furthermore, *in vivo* selection is necessarily highly stringent and recoveries at each round are exceedingly low. Conjugation further complicates the selection and may promote mutations that risk selecting undesired size variants (Supplemental Figure S4). Ultimately, we felt that assuring highly efficient conjugation was essential.

In light of our previous *in vivo* selections for GBM-specific unconjugated DNA aptamers (19), we wished to limit the number of variables changed in the present *in vivo* selections. We deliberately chose to use explants from the same GBM PDX model, G39, which is known to have a leaky blood brain barrier as previously discussed (19). Negative control ApDCs serve to probe the generic impact of this permeability. We find that *in vivo-*selected ApDCs accumulate in the orthotopic GBM at levels much higher than either negative control molecule. As negative controls, we opted to include Ap 7 as a control to compare unconjugated and conjugated products of *in vivo* SELEX, with the forward primer and random regions shared with negative control X (19). The increased retention of ApDC 6 vs. Ap 7 is hypothesized to reflect the presence of the hydrophobic cargo, and the existence of potential MMAE receptors in various tissues.

In the present *in vivo* SELEX experiment, an aptamer presumably was rewarded for importing its conjugated toxin cargo into the GBM. We hypothesize that the success of this SELEX was due in part to i) rigorous perfusion of the mouse to remove contaminating blood from tissues, ii) rapid tissue extraction and significant precautionary measures taken against cross-contamination during organ extraction, and iii) optimized extraction methodology that reproducibly recovered significantly more ApdC molecules. Notably, the percentage of the pool recovered as top candidates was significantly higher in this SELEX experiment than previously. Individual candidates, rather than clusters of similar candidates, could be evaluated.

Future studies will be needed to explore the mechanisms by which *in vivo* selected ApDCs home to the brain parenchyma and to GBMs. Although the *in vivo* selected ApDCs did not recognize mouse GBM or DMG tumors, the observation that trained ApDCs accumulated preferentially to untrained ApDCs in mouse brain is encouraging that the CNS homing phenomenon is not specific to the G39 model.

Although the GBM tumor is one of the primary tissues to which the *in vivo* selected ApDCs home, the overall biodistribution profile of the ApDCs is unexpected when compared to molecules selected in the absence of MMAE conjugation. It remains unclear if the observed lung accumulation is driven by the ApDC ligand or the presence of MMAE. Regardless, this finding illuminates a key limitation in the current *in vivo* SELEX strategy: the absence of a negative selection step. Given the cellular complexity in the composition of whole organs, we hypothesize that inclusion of a negative cell selection step may not be effective. However, methods to selectively eliminate lung-predominant sequences could be envisioned. Alternatively, development of a lung-decoy aptamer may also be a possible therapeutic approach.

Another important limitation of this work is that we did not develop methods to evaluate ApDC conjugation status *in vivo*. Because our quantitation method was based on highly-sensitive qPCR, measurements required the aptamer to be substantially intact and were blind to conjugation status. It is known that mouse serum carries carboxylesterase Ces1c, capable of cleaving the valine-citrulline linker of ADCs (37). Levels of injected ApDCs in our work would likely be below the limit of routine detection by LC-MS/MS. Given our finding that ApDCs lose target cell binding specificity in the absence of MMAE, we hypothesize that ApDCs from which MMAE has been prematurely cleaved *in vivo* may similarly lose the ability to home to the target GBM.

A unique challenge observed with *in vivo-*selected candidates from conjugated and unconjugated SELEX experiments is the inability to tolerate significant modifications post-SELEX. In a previous study using cell-selected candidates, one out of four selected aptamers tolerated conjugation to an antibody and successfully carried the antibody as cargo to its intracellular destination (38). Thus, the failure all *in vivo-*selected aptamers in two separate selections to tolerate cargo modifications requires further study, as it could be pointing to an underlying constraint required for aptamers to transit compartments *in vivo.* This also serves as a point of consideration for future *in vivo* SELEX experiments.

The described ApDC biodistribution and target tissue/cell binding are interesting, especially in the case of ApDC 1. This molecule is characterized by perhaps the most impressive anti-GBM biodistribution in this selection and subtly stains excised GBM tissue, yet there is no evidence of statistically significant binding to the cultured GBM cells. It is tempting to disregard this candidate, but it is possible that identifying the target of ApDC 1 may therapeutically relevant. If ApDC 1 binds to unique endothelial cells in the GBM, targeting tumor angiogenesis could be an effective therapy. Alternatively, if ApDC 1 recognizes host immune cells or fibroblasts, the molecule could be exploited for delivery of therapies to these specific cells.

Due to the selection criteria requiring ApDCs to survive in the tumor for four h post-injection, we note from the PK study that the selected ApDCs successfully accumulate and persist in the GBM. Thus, the *in vivo-*selected ApDCs have impressive biodistributions, anti-GBM homing, and pharmacokinetics. By 4h post-injection, the GBM tumor is receiving upwards of 50% of the remaining ApDC dose. From 0.5 to 8 h post-injection, ApDCs accumulate in the orthotopic tumors at levels comparable to those in well-perfused organs (e.g., lung and liver), a quality not observed in free MMAE or some ADCs (26, 39).

Herein, we present what we believe to be a first example of *in vivo* SELEX involving toxin-conjugated DNA library. Four of the five top ApDCs from this selection reproducibly bind the GBM tumor *in vivo*, preserved GBM tissue *ex vivo*, and the target GBM cells in culture. Besides demonstrating *in vivo* GBM homing preference we detect surprising accumulation in the lung as has been reported for both free MMAE and for ADCs (26, 40). This work provides a first demonstration of an effective method for developing anti-GBM homing ApDCs. Future studies will explore the tumor toxicity of the selected reagents.

## Author Contributions

CDD, SJ, BAW, DMB, JNS and LJM conceived and designed the experiments. CDD, SJ, LLO, and KKB performed the experiments. KSP, JG, ZT, JAM, and SY performed an experiment. CDD performed data analysis, CDD and LJM wrote the manuscript.

## Acknowledgments

This work was supported by Mayo Clinic (L.J.M., J.N.S); L.J.M. is the Bernard Pollack Professor and J.N.S. is the William H. Donner Professor at Mayo Clinic. Work also supported by NIH grants T32GM145408 Mayo Clinic Medical Scientist Training Program and F30CA294722-01A1 (C.D.), K99NS138490 (S.Y.), R35GM143949 (L.J.M.), R61NS128071-01A1 (J.S., L.J.M., W.F.E.), Mayo Clinic CTSA grant UL1TR000135, a Mayo Clinic CCaTS-CBD Team Science award, a generous grant from the Humor to Fight the Tumor Foundation, and an NSF graduate fellowship (B.W.).HLF and 3T3 cells were generously given by A. Haak, PhD at Mayo Clinic. H3K27MPP cells were generously provided by Dr. Sameer Agnihotri at the University of Pittsburgh. The graphical abstract, Figure 2A, and Supplemental Figure S5 were created with BioRender.

## Data sharing

All data are provided in supplemental materials. Constructs are available from the authors upon request.

