## Supplemental files for "*In Vivo* Selection of anti-glioblastoma DNA aptamer-drug conjugates in an orthotopic patient-derived xenograft model"

### Table of Contents

### 1.0 Supplemental data

**Supplemental Table 1**. Oligonucleotides used in experiments.
**Supplemental Table 2.** qPCR values of sequence specific primers.
**Supplemental Table 3**. Evaluation of cross-reactivity between candidates and sequence- specific primers.
**Supplemental Table 4.** Mice used for in vivo SELEX selections.
**Supplemental Figure S1.** Binding of aptamers selected *in vivo* with an unconjugated library to target cells (G39) detected by qPCR
**Supplemental Figure S2.** Rationale for transition from trans-cyclooctene-tetrazine (TCO-Tz) click chemistry to commercially available 5'MMAE-conjugated primers.
**Supplemental Figure S3.** Evaluating MMAE for compatibility with SELEX conditions.
**Supplemental Figure S4**. Library length variation in initial selection.
**Supplemental Figure S5.** Diagram of toggle "crick" approach.
**Supplemental Figure S6.** Evaluation of libraries after SELEX.
**Supplemental Figure S7.** qPCR analysis of ApDC library during selection rounds and in the indicated organs of the round 10 mouse
**Supplemental Figure S8.** ApDC accumulation in lung is observed from in vivo selected candidates with a conjugated library.
**Supplemental Figure S9.** Evaluating the library more deeply for primarily tumor homing aptamers.
**Supplemental Figure S10.** ApDC tissue selectivity is observed by post-staining regardless of method: OCT or FFPE.
**Supplemental Figure S11.** Additional *ex vivo* tissue staining.
**Supplemental Figure S12.** *Ex vivo* tissue staining on brain sections from the mouse shown in Supplemental Figure S14 (Mouse 9037), with tumor sections removed from the tissue.
**Supplemental Figure S13.** *Ex vivo* tissue staining on brain sections from the same mouse as in Supplemental Figure S13 (Mouse 9037) with tumor.
**Supplemental Figure S14.** Additional ApDC binding to target cells in culture.
**Supplemental Figure S15.** Predicted secondary structures for the indicated aptamers as determined using mFold

**Supplemental Figure S16.** Evaluating human G39 selected ApDC specificity to mouse high grade tumor models

### 2.0 Supplemental references

### 1.0 Supplemental Data

**Supplemental Table S1. Experimental oligonucleotides.**

| Internal Name | Description | Sequence |
|---|---|---|
| LJM-7166 | 5' MMAE forward SELEX primer | /5VCPMPEG4N/AGTCTGTTCTCCTGTCTCAG |
| LJM-7468 | *In vivo* SELEX library | AGTCTGTTCTCCTGTCTCAG($N_{40}$) GCGGATCTAAGACATCTAGC |
| LJM-7542 | Reverse SELEX primer (56nt) | /56-FAM/AAAAAAAAAAAAAAAAAAAAAAAAAAAAAAAAAAAA/iSp9//iSp9/GCTAGATGTCTTAGATCCGC |
| LJM-7541 | Reverse SELEX primer 2 (20nt) | /56-FAM/GCTAGATGTCTTAGATCCGC |
| LJM-7540 | Reverse SELEX primer 3 (50nt) | /56-FAM/AAAAAAAAAAAAAAAAAAAAAAAAAAAAAA/iSp9//iSp9/GCTAGATGTCTTAGATCCGC |
| LJM-6880 | Unmodified forward primer | AGTCTGTTCTCCTGTCTCAG |
| LJM-7470 | Unmodified reverse primer | GCTAGATGTCTTAGATCCGC |
| LJM-7589 | ApDC 1 | **AGTCTGTTCTCCTGTCTCAG**GCGCAAGCCGGTATTTGGTGCTAAAGTCTATCAAGTCCTC **GCGGATCTAAGACATCTAGC** |
| LJM-7591 | ApDC 2 | **AGTCTGTTCTCCTGTCTCAG**TTAACCCCTCACATCTCAATCCTTCTTTCGTCGTTCCCTC **GCGGATCTAAGACATCTAGC** |
| LJM-7593 | ApDC 3 | **AGTCTGTTCTCCTGTCTCAG**TTACAATCACCCCGACCTCTCTCTTATTGTTGTCCTGTTA **GCGGATCTAAGACATCTAGC** |
| LJM-7595 | ApDC 4 | **AGTCTGTTCTCCTGTCTCAG**TAGTTGCTAAATCCCGTTTTCATCTTCTATTCTTGTCCTC **GCGGATCTAAGACATCTAGC** |
| LJM-7597 | ApDC 5 | **AGTCTGTTCTCCTGTCTCAG**CGATAGCCACTTGCTTTAAACCCTCCTTTCTTGAACGGTA **GCGGATCTAAGACATCTAGC** |
| LJM-7599 | ApDC 6 | **AGTCTGTTCTCCTGTCTCAG**CGAAGATCATTTTCTTGTTACTCGCGGGTCACAGAACGCG **GCGGATCTAAGACATCTAGC** |
| LJM-7601 | Primer for ApDC 1 | GAGGACTTGATAGACTTTAG |
| LJM-7602 | Primer for ApDC 2 | GAACGACGAAAGAAGG |
| LJM-7603 | Primer for ApDC 3 | GACAACAATAAGAGAGAGGT |
| LJM-7604 | Primer for ApDC 4 | CAAGAATAGAAGATGAAAAC |
| LJM-7605 | Primer for ApDC 5 | GTTCAAGAAAGGAGGG |
| LJM-7606 | Primer for ApDC 6 | GCGTTCTGTGACCCG |

**Supplemental Table S2. qPCR values of sequence specific primers.**

| | Slope | Intercept |
|---|---|---|
| **ApDC 1** | -2.78 | 15.78 |
| **ApDC 2** | -1.56 | 12.06 |
| **ApDC 3** | -2.75 | 15.70 |
| **ApDC 4** | -2.80 | 15.83 |
| **ApDC 5** | -2.90 | 15.90 |
| **ApDC6** | -2.85 | 16.70 |

**Supplemental Table S3. Evaluation of cross-reactivity between candidates and sequence-specific primers.** Average $C_q$ values obtained from testing the sequence specific primers (horizontal) against the various sequences (1nM, vertical); primer specificity ($P_s$) were $L_B$ is the $C_q$ value for library A amplified with library B primers and $L_A$ is the $C_q$ value for Library A amplified with Library A primers.

**Averages**

| | ApDC_1 (7589) | ApDC_2 (7591) | ApDC_3 (7593) | ApDC_4 (7595) | ApDC_5 (7597) | ApDC_6 (7599) |
|---|---|---|---|---|---|---|
| ApDC_1 (7589) | 7.232052742 | 24.3842264 | 25.98216575 | 32.60932775 | 32.78963872 | 30.19095056 |
| ApDC_2 (7591) | 16.8475858 | 7.685762948 | 22.79255217 | 32.75439638 | 28.85452684 | 32.04276891 |
| ApDC_3 (7593) | 28.47338718 | 24.46955143 | 6.877157802 | 29.67978729 | 29.25734752 | 29.09279677 |
| ApDC_4 (7595) | 23.68825385 | 23.69852169 | 23.69716981 | 7.246881004 | 22.75988499 | 23.41812244 |
| ApDC_5 (7597) | 0 | 0 | 0 | 0 | 23.25565381 | 0 |
| ApDC_6 (7599) | 34.90819484 | 34.16940293 | 34.24304425 | 32.6713116 | 25.67510775 | 7.964444926 |

**Calculation Ps = 2^(LB-LA)**

| | ApDC_1 (7589) | ApDC_2 (7591) | ApDC_3 (7593) | ApDC_4 (7595) | ApDC_5 (7597) | ApDC_6 (7599) |
|---|---|---|---|---|---|---|
| ApDC_1 (7589) | 1 | 145652.774 | 440906.436 | 43583387.82 | 49385561.34 | 8152990.004 |
| ApDC_2 (7591) | 572.7742617 | 1 | 35285.53404 | 35189298.84 | 2357397.948 | 21487673.64 |
| ApDC_3 (7593) | 3170391.076 | 197623.3142 | 1 | 7316029.656 | 5458941.166 | 4870501.739 |
| ApDC_4 (7595) | 88991.06355 | 89626.6829 | 89542.73749 | 1 | 46760.54037 | 73795.30624 |
| ApDC_5 (7597) | 9.98506E-08 | 9.98506E-08 | 9.98506E-08 | 9.98506E-08 | 1 | 9.98506E-08 |
| ApDC_6 (7599) | 129085339.3 | 77353219.8 | 81404167.56 | 27384698.67 | 214506.6463 | 1 |

**Check if >100,000**

| | ApDC_1 (7589) | ApDC_2 (7591) | ApDC_3 (7593) | ApDC_4 (7595) | ApDC_5 (7597) | ApDC_6 (7599) |
|---|---|---|---|---|---|---|
| ApDC_1 (7589) | FALSE | 0 | 0 | 0 | 0 | 0 |
| ApDC_2 (7591) | FALSE | FALSE | FALSE | 0 | 0 | 0 |
| ApDC_3 (7593) | 0 | 0 | FALSE | 0 | 0 | 0 |
| ApDC_4 (7595) | FALSE | FALSE | FALSE | FALSE | FALSE | FALSE |
| ApDC_5 (7597) | FALSE | FALSE | FALSE | FALSE | FALSE | FALSE |
| ApDC_6 (7599) | 0 | 0 | 0 | 0 | 0 | FALSE |

| Cohort 1 (04-25-24 injection) | | | | | | |
|---|---|---|---|---|---|---|
| **Cell Density** | **Round** | **Mouse ID** | **Collected (Study Day)** | **Peak BLI value (day)*** | **Final BLI value (day)** | **IC Tumor Weight (mg)** |
| **100k** | 1 | 2531 | 5-14-24 (19) | | 5-13-24 (18) 2.07 E+09 | 145.6 |
| | ~~2~~ | ~~6180~~ | ~~5-16-24 (21)~~ | | ~~5-16-24 (21) 3.22 E+09~~ | ~~128.5~~ |
| | ~~3~~ | ~~7406~~ | ~~5-17-24 (22)~~ | | ~~5-16-24 (21) 1.39 E+09~~ | ~~81.3 (hemorrhage)~~ |
| **10k** | Repeat 2 | 6835 | 5-19-24 (24) | | 5-19-24 (24) 8.45 E+08 | 78 |
| | 3.1 | 4247 | 5-21-24 (26) | | 5-21-24 (26) 6.28 E+08 | 123.9 |
| | 4 | 4057 | 5-24-24 (29) | | 5-23-24 (28) 8.42 E+08 | 144.6 |
| | 5 | 1619 | 5-25-24 (30) | 5-23-24 (28) 7.90 E+08 | 5-25-24 (30) 5.19 E+07 | 74.8 |
| **1k** | ~~6~~ | ~~1990~~ | ~~5-27-24 (32)~~ | ~~5-25-24 (30) 1.19 E+09~~ | ~~5-27-24 (32) 9.26 E+08~~ | ~~86.6 (hemorrhage)~~ |
| | Repeat 6 | 3006 | 5-28-24 (33) | | 5-27-25 (32) 9.38 E+08 | 117.6 |
| | 7 | 4836 | 5-31-24 (36) | 5-27-24 (32) 9.03 E+08 | 5-30-24 (35) 6.89 E+08 | 109.1 |
| | 8 | 1692 | 6-3-24 (39) | | 5-30-24 (35) 1.41 E+08 | 32.7 |

| Cohort 2 (05-23-24 injection) | | | | | | |
|---|---|---|---|---|---|---|
| **Cell Density** | **Round** | **Mouse ID** | **Collected (Study Day)** | **Peak BLI value (day) *** | **Final BLI value (day)** | **IC Tumor Weight (mg)** |
| **100k** | 9 | 7156 | 6-7-24 (15) | | 6-7-24 (15) 5.46 E+08 | 23.9 |
| | 10 | 6106 | 6-10-24 (18) | | 6-10-24 (18) 6.29 E+08 | 16.8 |
| **50K** | 4A | 4470 | 6-20-24 (28) | 6-17-24 (25) 1.48 E+09 | 06-20-24 (28) 8.95 E +07 | 113 |
| | 5A | 3812 | 6-24-24 (32) | | 6-24-24 (32) 6.94 E+09 | 89.4 |
| **10k** | 6A | 4798 | 6-26-24 (34) | | 6-26-24 (34) 2.14 E+08 | 23.4 |
| | 7A | 2826 | 7-1-24 (39) | | 6-26-24 (34) 2.04 E+08 | 9.6 |
| | 8A | 1024 | 7-5-24 (43) | 6-24-24 (32) 4.44 E+07 | 7-2-24 (40) 3.16 E+07 | 20.7 |
| **1k** | 9A | 5037 | 7-18-24 (56) | | 7-18-24 (56) 8.52 E+08 | 78.1 |
| | 10A | 3672 | 7-23-24 (63) | | 7-23-24 (63) 3.27 E+08 | 40.8 |
| | 11A | 6411 | | | | |
| | 12A | 3065 | | | | |

**Supplemental Table S4. Mice used for *in vivo* SELEX selections.** Log flux indicates bioluminescence imaging (BLI) indicating the presence of luciferase tagged PDX GBM tumor. Tumor masses do not correlate well with BLI readings.

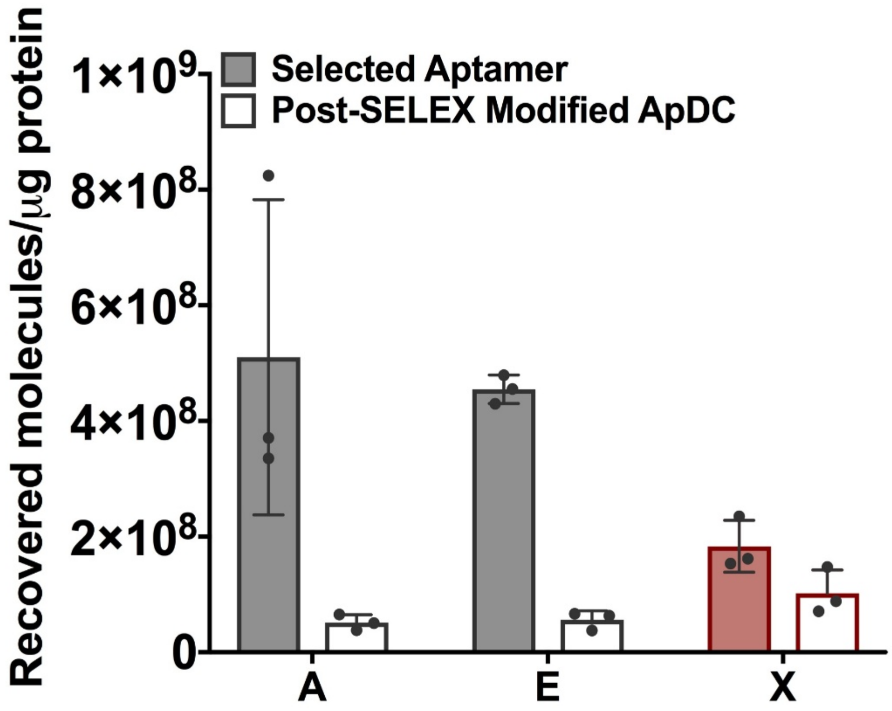


**Supplemental Figure S1.** Binding to target cells (G39) detected by qPCR. These aptamers were selected as unconjugated molecules as described in Doherty, et al., 2025 (1). In vitro binding was measured by qPCR for aptamers in their selected state (unconjugated) or after post-SELEX conjugation to MMAE (ApDCs created with 5'MMAE conjugated primers via the standard PCR method).

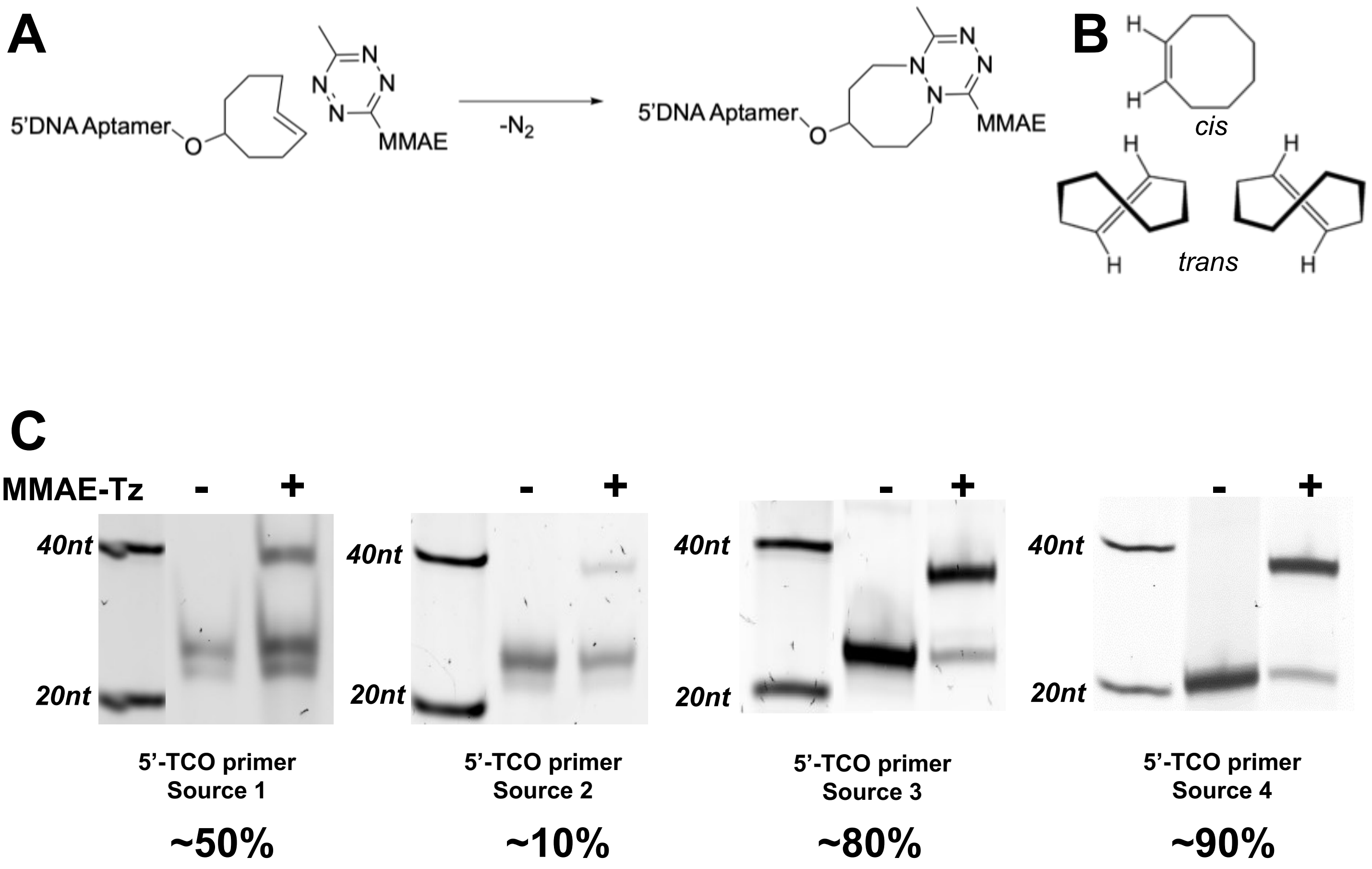


**Supplemental Figure S2. Rationale for transition from trans-cyclooctene-tetrazine (TCO-Tz) click chemistry to commercially available (MMAE-NHS ester and 5'-amino DNA oligonucleotides followed by HPLC purification ) 5'MMAE-conjugated primers.** A. A schematic demonstrating the isomerization that can occur with TCO to cis-cyclooctene, rendering the molecule unreactive. B. Polyacrylamide denaturing gels demonstrate the high variability in reactivity between batches of 5'TCO primers.

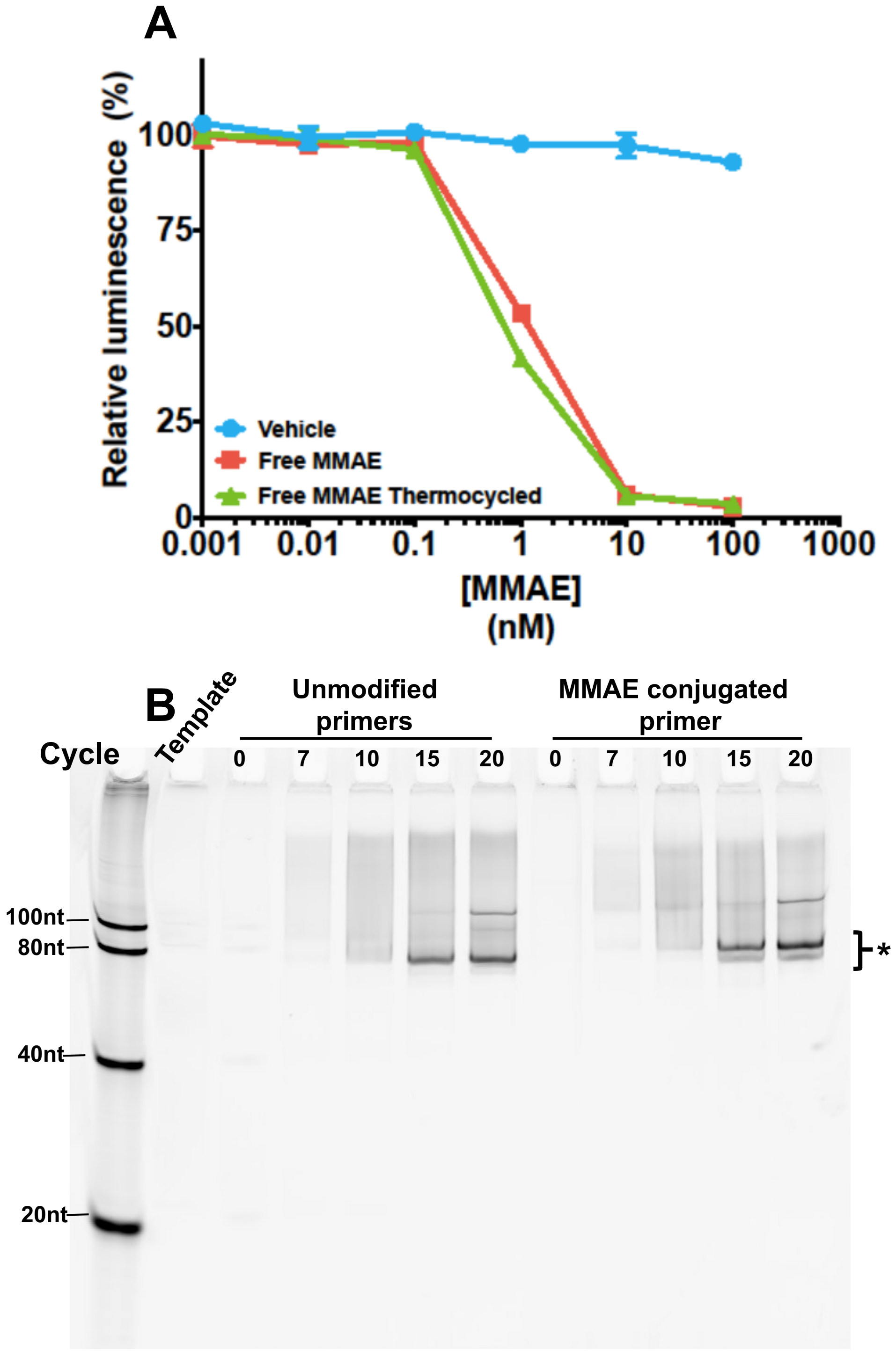


**Supplemental Figure S3. Evaluating MMAE for compatibility with SELEX conditions.** A. Toxicity of free MMAE to the target line G39 with or without 25 cycles of thermocycling. B. Same gel shown in Fig. 1D imaged with SYBR green (detects only duplexes created during PCR) prior to staining with SYBR gold.

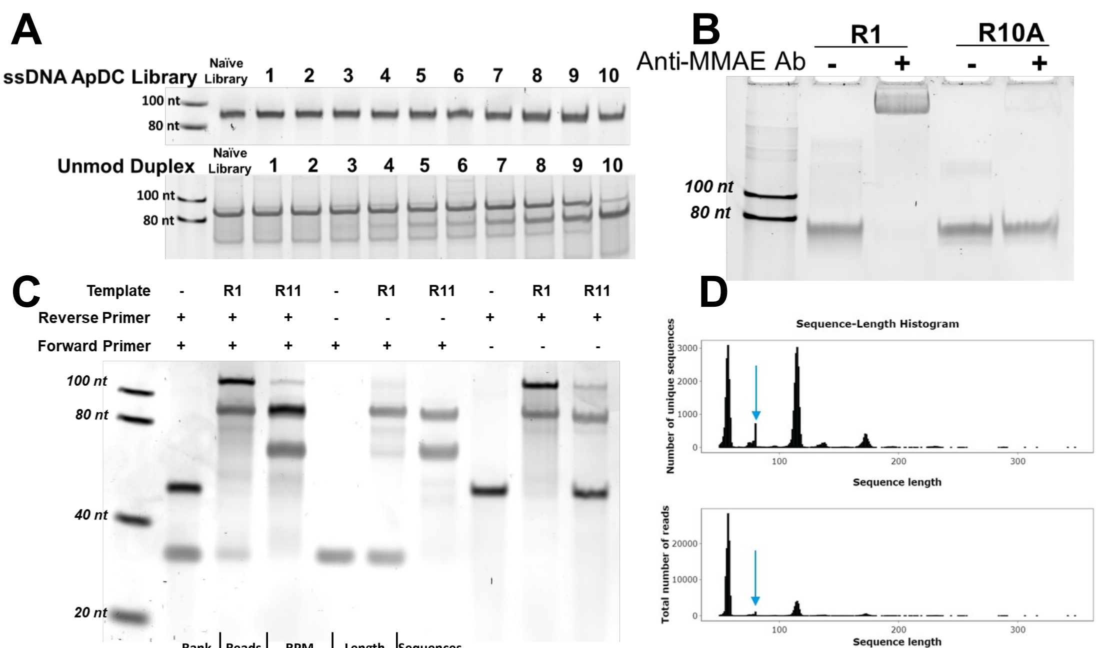


E

| Rank | Reads | RPM | Length | Sequences |
|---|---|---|---|---|
| 1 | 2673 | 23805.49 | **58** | AGTCTGTTCTCCTGTCTCAGAGAGAGCTTCCCTGACCAGCGGATCTAAGACATCTAGC |
| 3 | 1787 | 15914.86 | **57** | AGTCTGTTCTCCTGTCTCAGGCTCTGCCTCGAGGGAAGCGGATCTAAGACATCTAGC |
| 5 | 1097 | 9769.78 | **57** | AGTCTGTTCTCCTGTCTCAGGTTGGGATAATACCGTGGCGGATCTAAGACATCTAGC |
| 7 | 1052 | 9369.02 | **55** | AGTCTGTTCTCCTGTCTCAGGCTGTCGATCTCCCTGCGGATCTAAGACATCTAGC |
| 9 | 877 | 7810.48 | **57** | AGTCTGTTCTCCTGTCTCAGAAAGAGGGGTTGGCAGTGCGGATCTAAGACATCTAGC |
| 11 | 826 | 7356.28 | **55** | AGTCTGTTCTCCTGTCTCAGTAAGGGGCAAGTGGTGCGGATCTAAGACATCTAGC |
| 13 | 738 | 6572.56 | **59** | AGTCTGTTCTCCTGTCTCAGCCATTGGAGTGCTGGTTTTGCGGATCTAAGACATCTAGC |
| 15 | 733 | 6528.03 | **57** | AGTCTGTTCTCCTGTCTCAGGACACGACTATCCCGCTGCGGATCTAAGACATCTAGC |
| 17 | 695 | 6189.61 | **56** | AGTCTGTTCTCCTGTCTCAGCCTCCCAAGTAGACTAGCGGATCTAAGACATCTAGC |
| 19 | 611 | 5441.51 | **56** | AGTCTGTTCTCCTGTCTCAGGAGATGGCTGAGGTGGGCGGATCTAAGACATCTAGC |

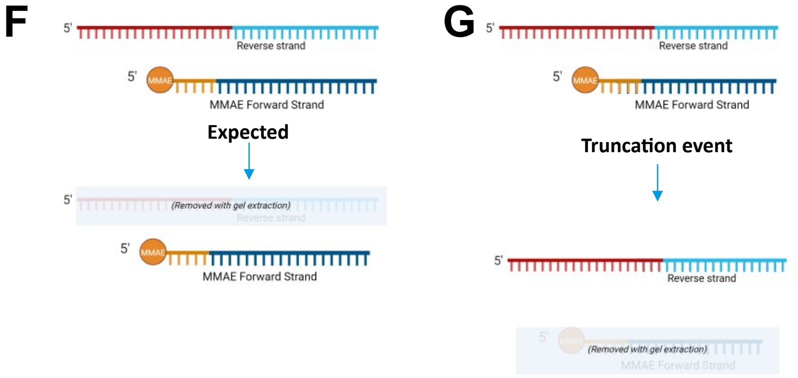


**Supplemental Figure S4. Library length variation in initial selection.** The first SELEX attempt involved only one reverse primer (20 nt) for all cycles. Length variation mutations were common enough that library member lacking MMAE migrated at the position expected for MMAE conjugates, preventing proper product purification at each cycle. This was avoided by alternating reverse primers of three sizes to facilitate purification of the proper MMAE conjugate library. A. Denaturing gel of 5'MMAE libraries over the 10 rounds of the initial selection attempt. Libraries migrate near the 90-nt marker . B. Native gel of unconjugated duplexes over the 10 rounds of selection. By Round 4 a spurious, smaller duplex begins to appear, which dominates the library by Round 10. C. Native gel evaluating MMAE conjugation using an anti-MMAE antibody. D. PCR test to evaluate if the forward and/or reverse strand is present in the library at Round 10. E. Sequencing data confirming spurious library sequences whose mobility is comparable to 5'MMAE ApDC on a denaturing gel. F. Top 10 sequences in the library at Round 10. G-H. Schematic diagrams showing expectation (G) versus the apparent truncation product (H) resulting in a library lacking MMAE conjugation.

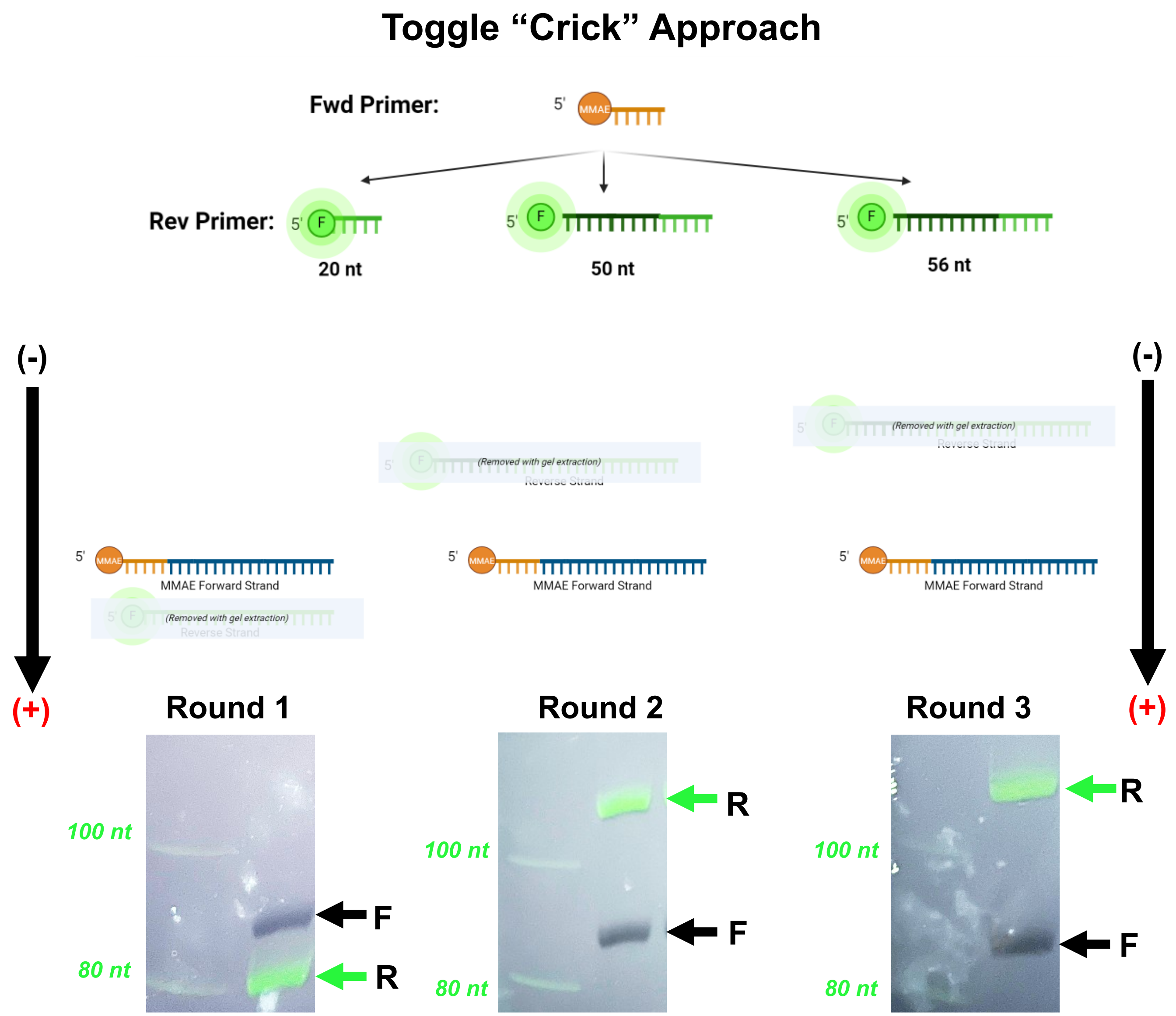


**Supplemental Figure S5.** Schematic diagram showing the reverse primer toggling strategy to facilitate proper 5'MMAE strand purification in the presence of aptamer length variants.

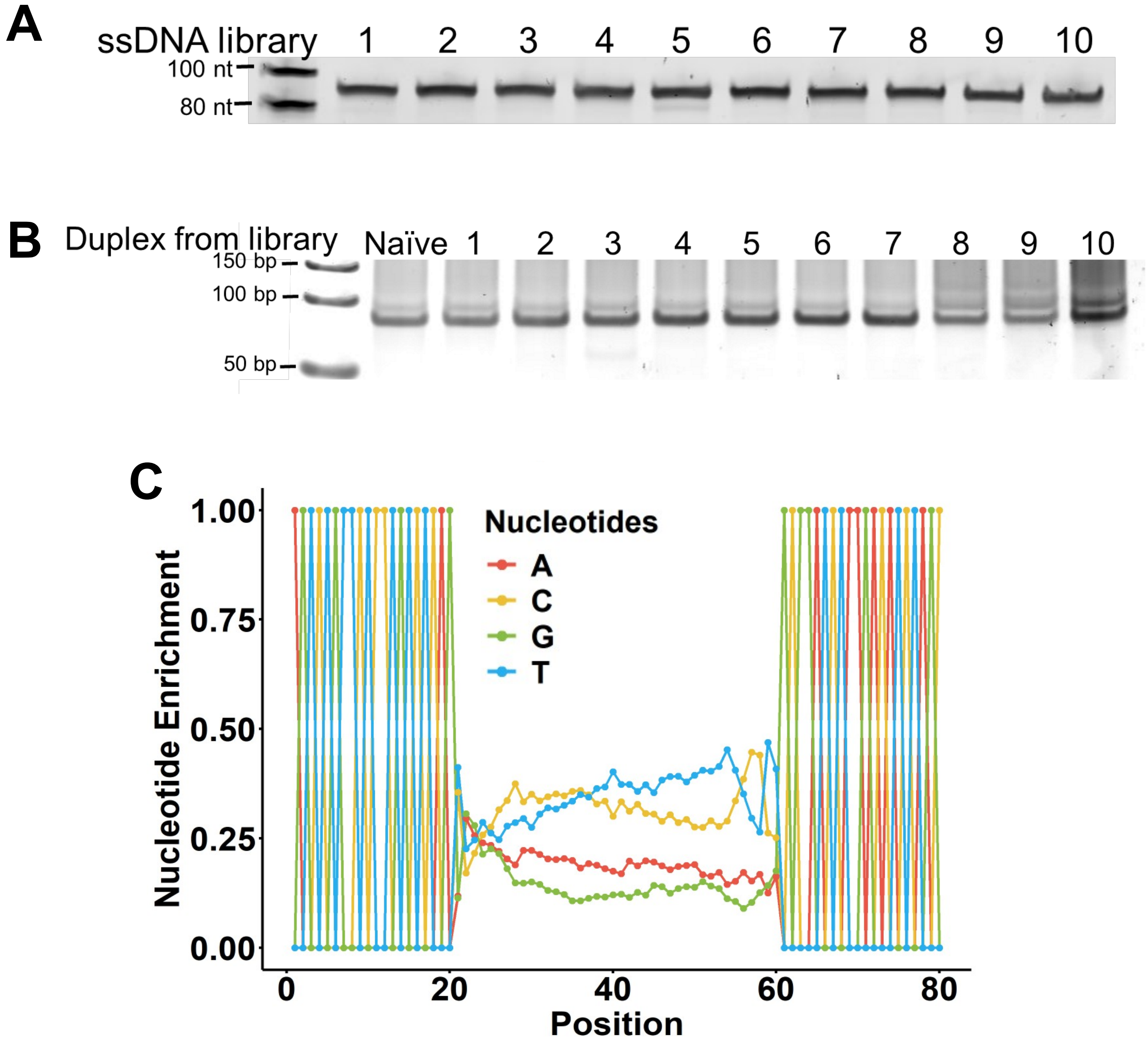


**Supplemental Figure S6. Evaluation of libraries after SELEX.** A. Denaturing gel of libraries demonstrating MMAE conjugation. B. Native polyacrylamide gel of the duplexes demonstrating that apparent library length of 80 bp. C. Nt distribution in the library at Round 10, from deep sequencing.

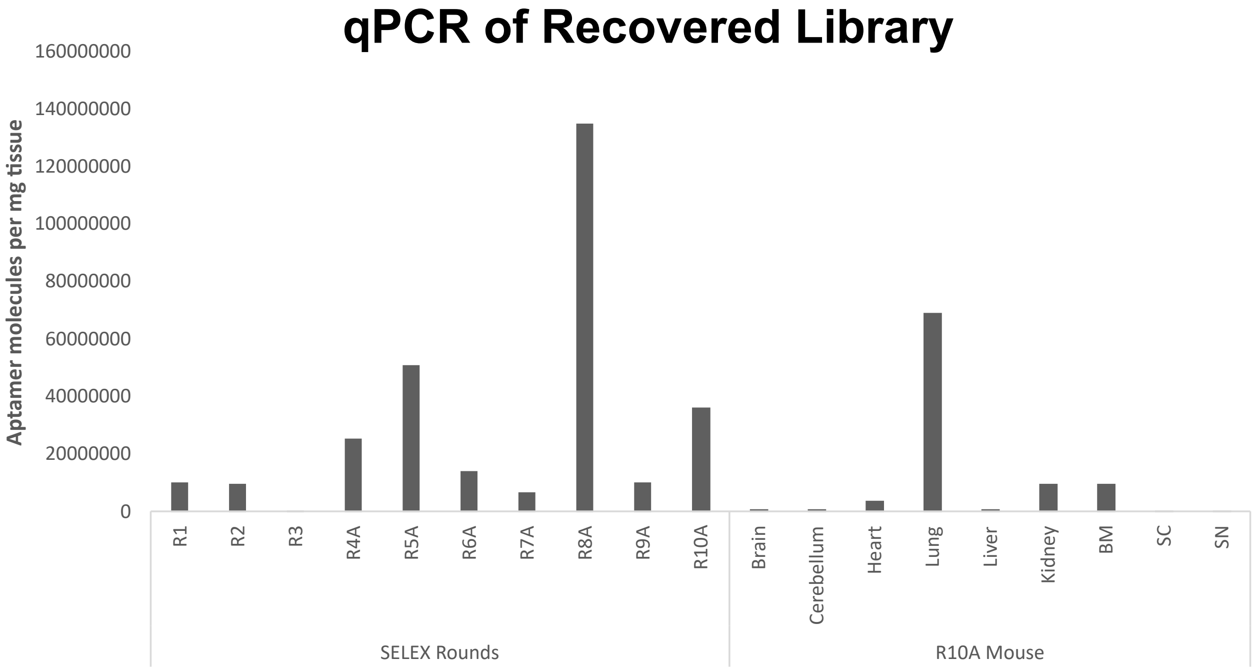


**Supplemental Figure S7.** qPCR analysis of ApDC library during selection rounds and in the indicated organs of the Round 10 mouse

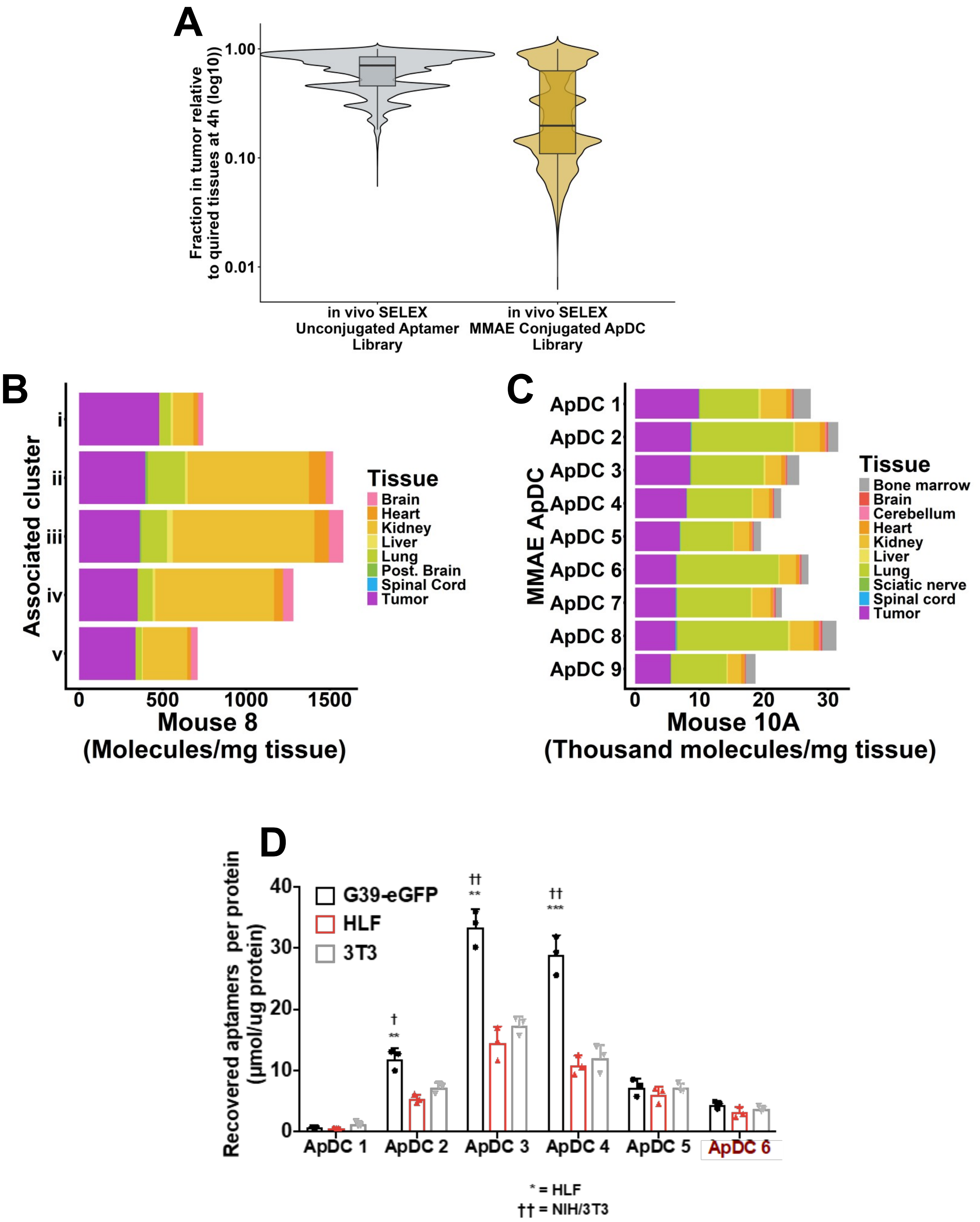


**Supplemental Figure S8. ApDC accumulation in lung is observed from *in vivo*-selected candidates with a conjugated library.** A. Comparison of the fraction of recovered injected dose at 4 h post-injection of the libraries from present *in vivo* SELEX vs. results for an unconjugated aptamer library (Doherty et al., 2025). B. Biodistribution of the top 5 clusters in the Round 8 mouse from previous *in vivo* SELEX with unconjugated molecules (Doherty et al., 2025). C. Biodistribution in the Round 10 mouse for the top 9 sequences. D. Cell binding to human (HLF) and mouse (NIH/3T3) lung fibroblasts.

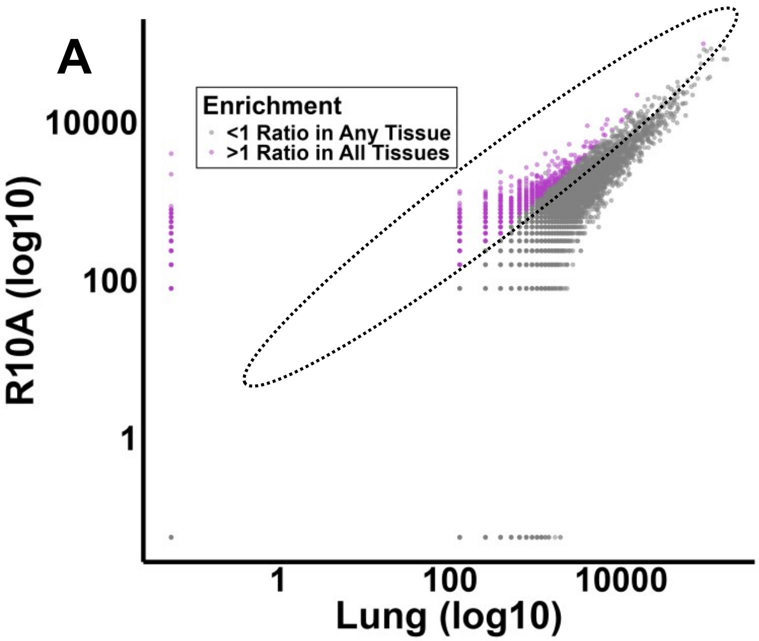


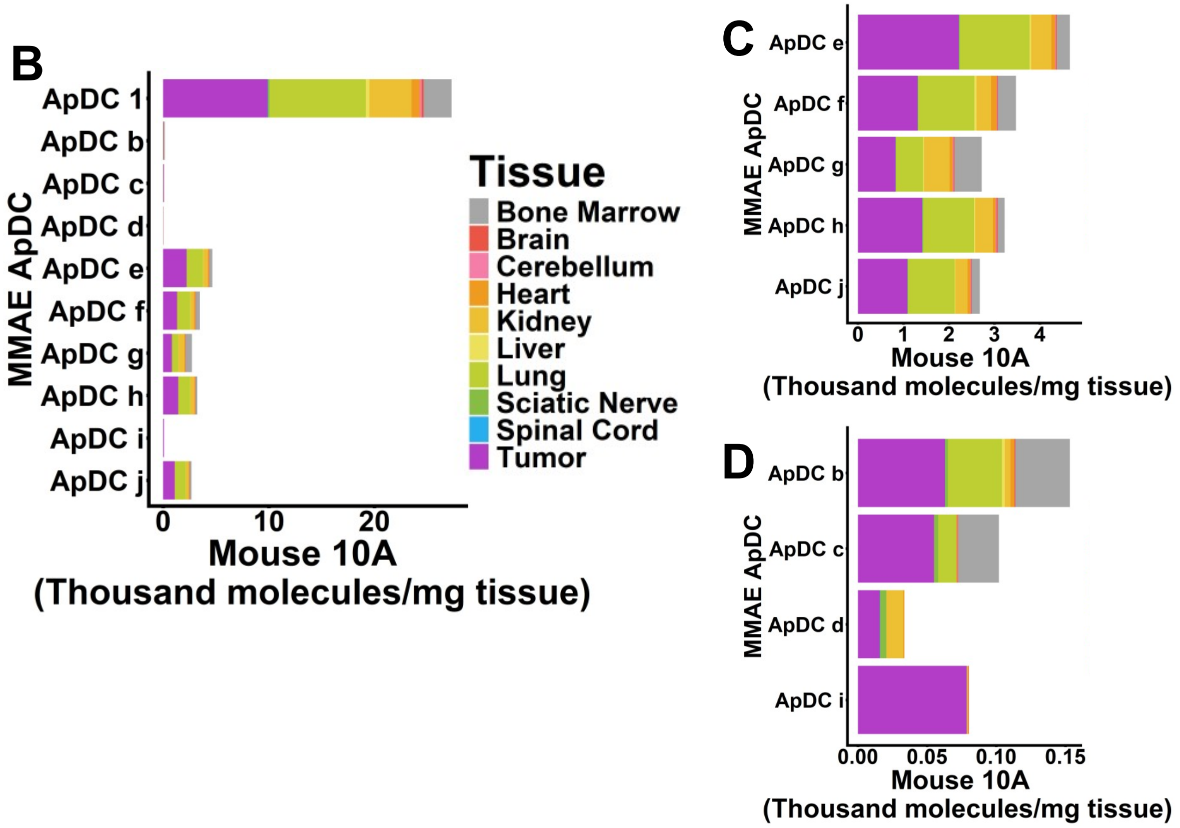


**Supplemental Figure S9. Evaluating the library more deeply for primarily tumor homing aptamers**. A. All 80-mers in the library evaluated for ratio of molecules found in the tumor relative to any other organ. B-D. Biodistribution of the top 10 aptamers that had tumor: any tissue ratios >1 (B). Other than ApDC 1, results further divided into ApDCs with total aptamer accumulation less than 5,000 molecules/mg tissue (C) and less than 150 molecules/mg tissue (D).

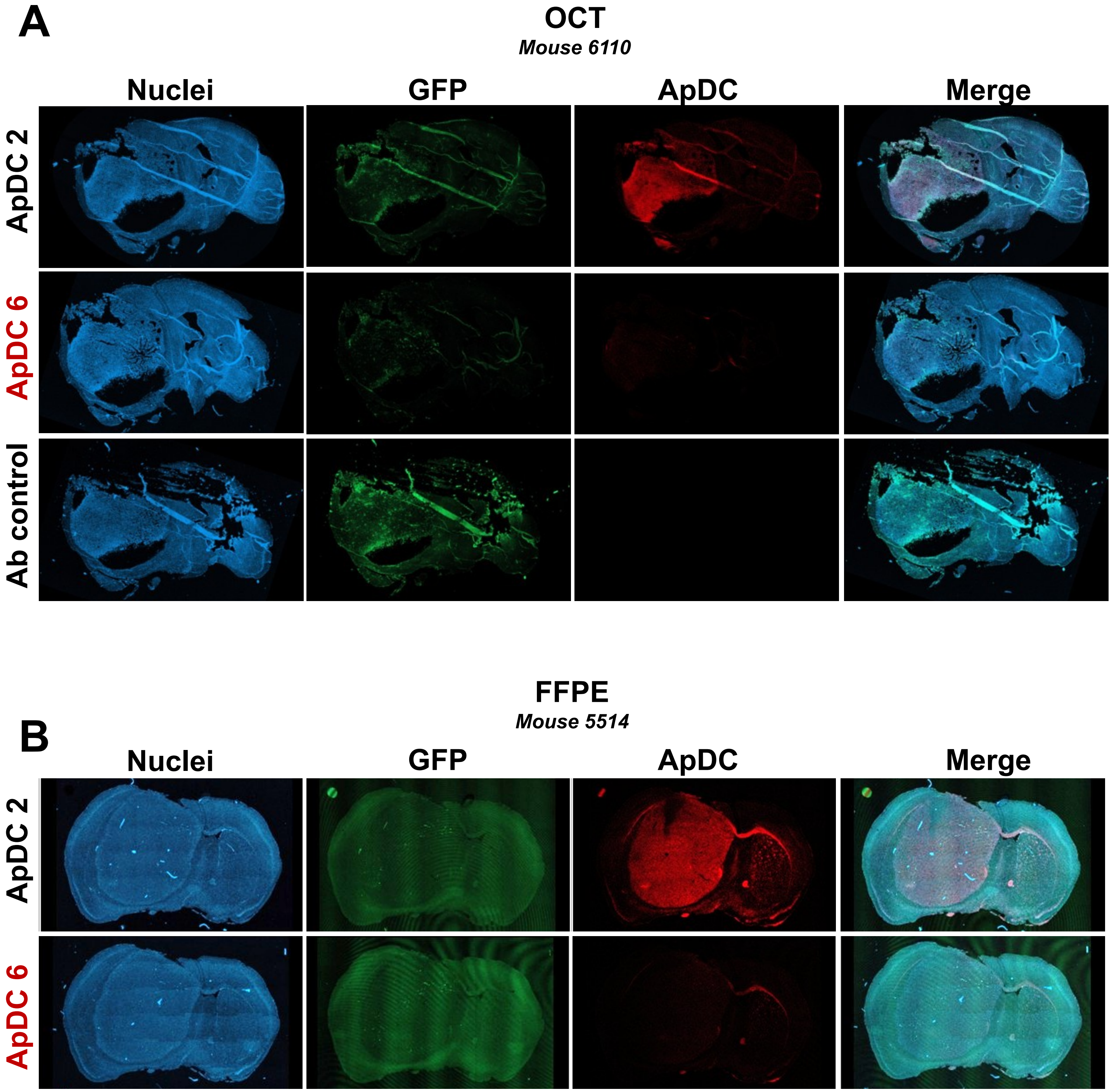


**Supplemental Figure S10.** ApDC tissue selectivity is observed by post-staining regardless of method: OCT (A) or FFPE (B).

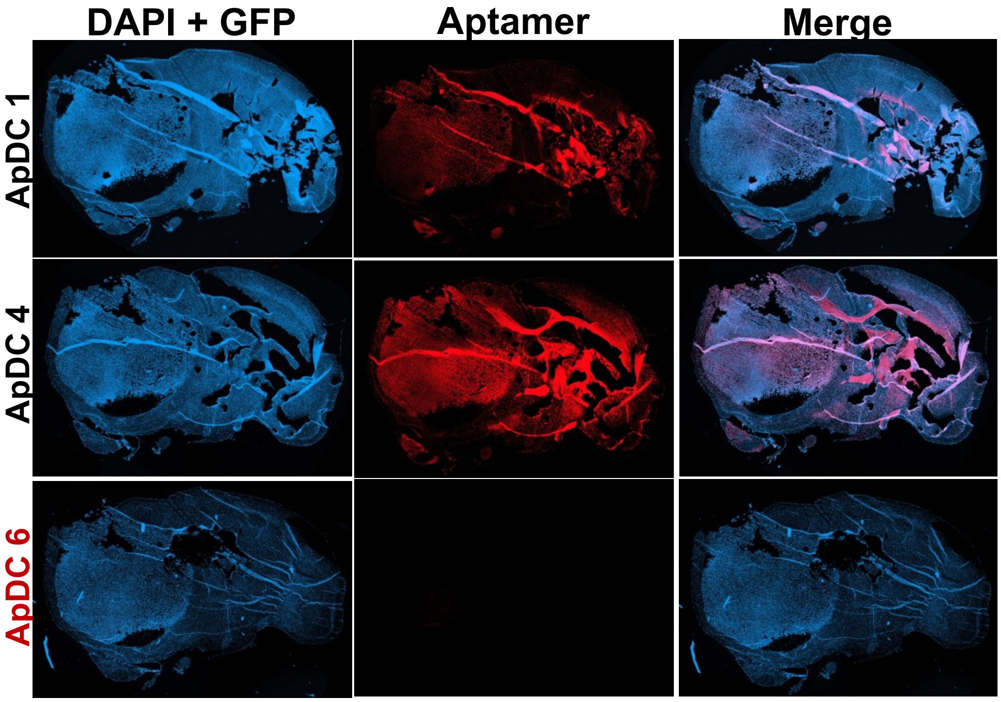


**Supplemental Figure S11.** *Ex vivo* tissue staining of indicated candidate ApDCs on serial brain sections from the same mouse as shown in Fig. 3.

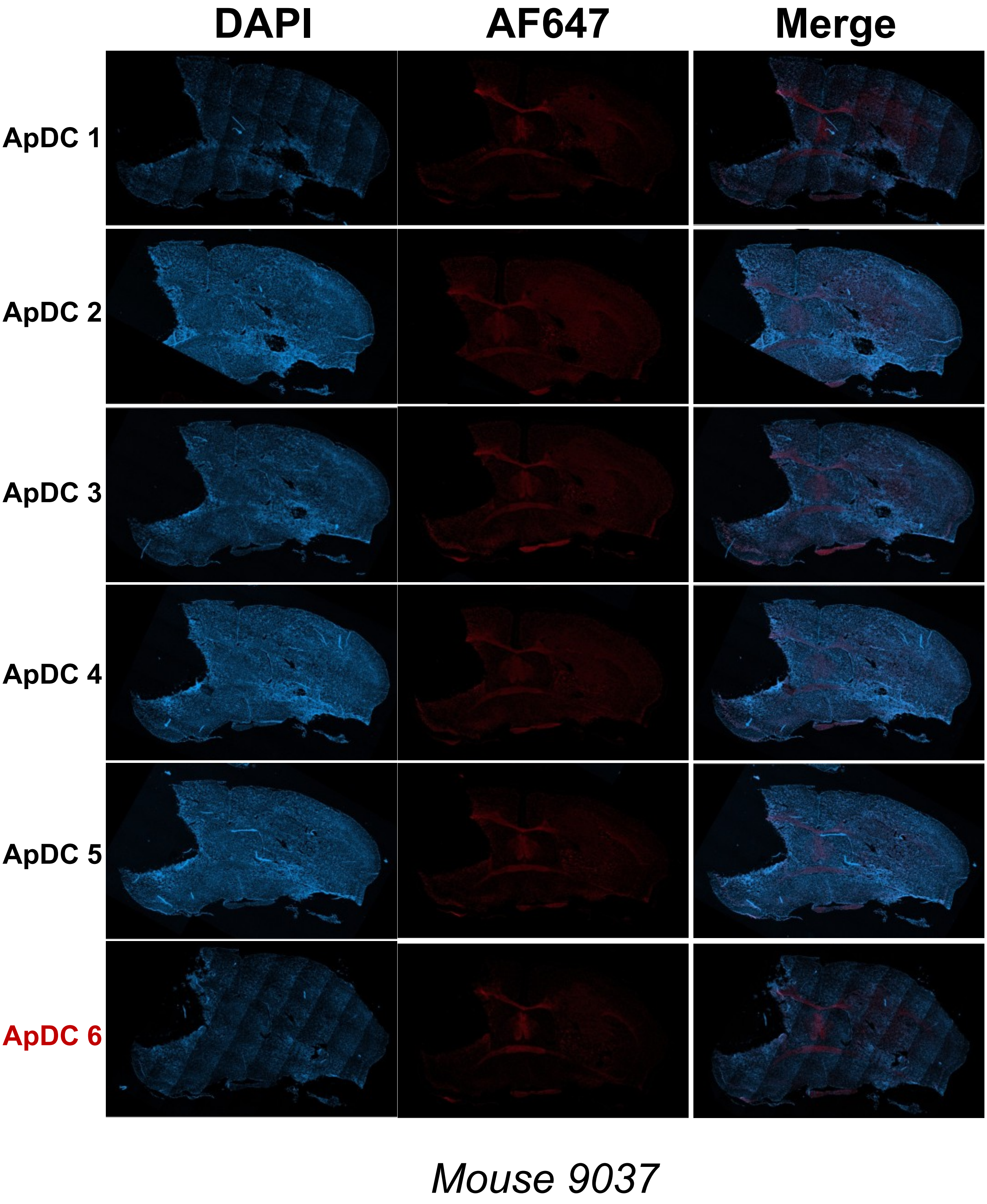


**Supplemental Figure S12.** *Ex vivo* tissue staining on brain sections from the mouse shown in Supplemental Figure S14 (Mouse 9037), with tumor sections removed from the tissue.

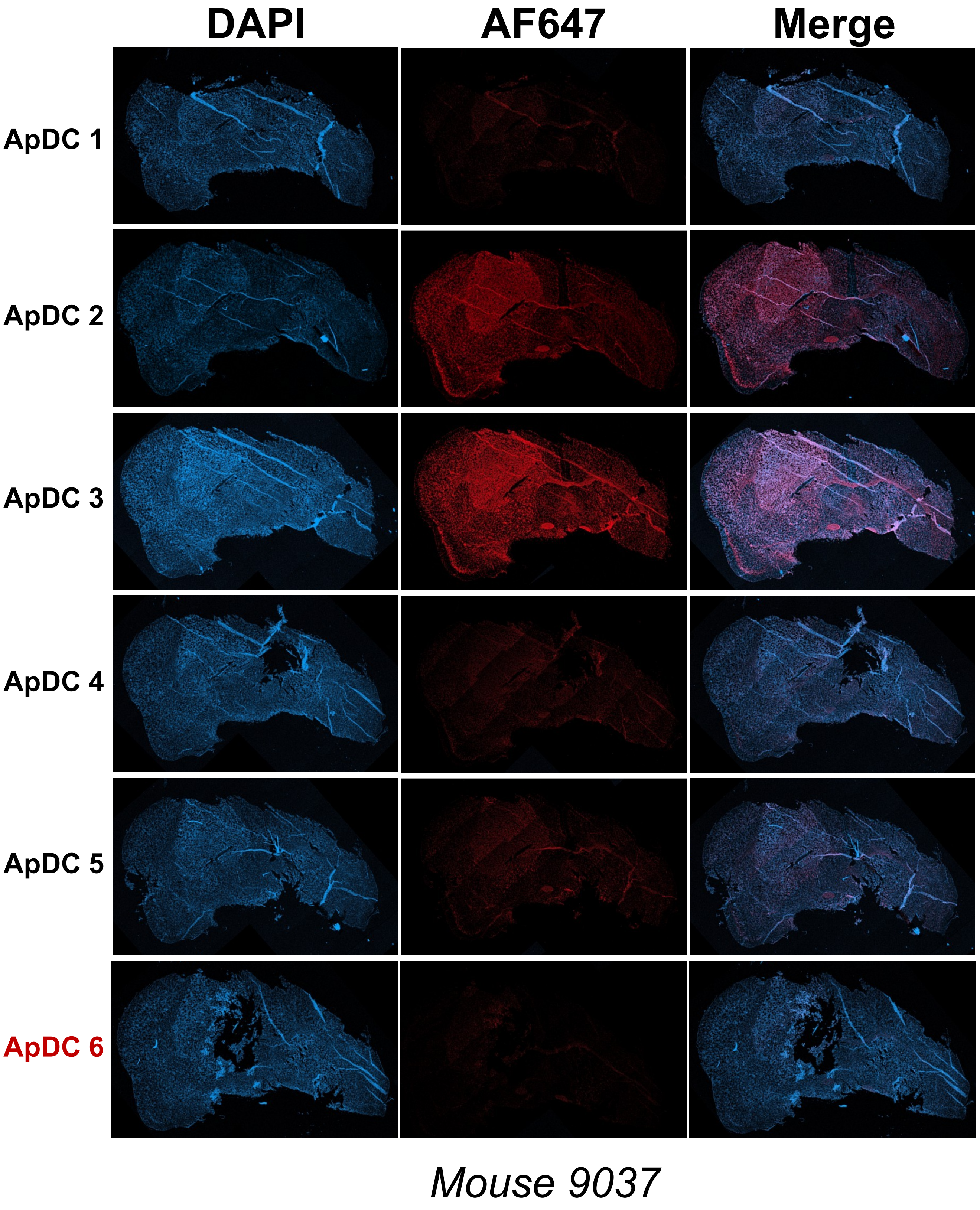


**Supplemental Figure S13.** *Ex vivo* tissue staining on brain sections from the same mouse as in Supplemental Figure S13 (Mouse 9037). Tissue sections are in posterior parts of the brain and tumor.

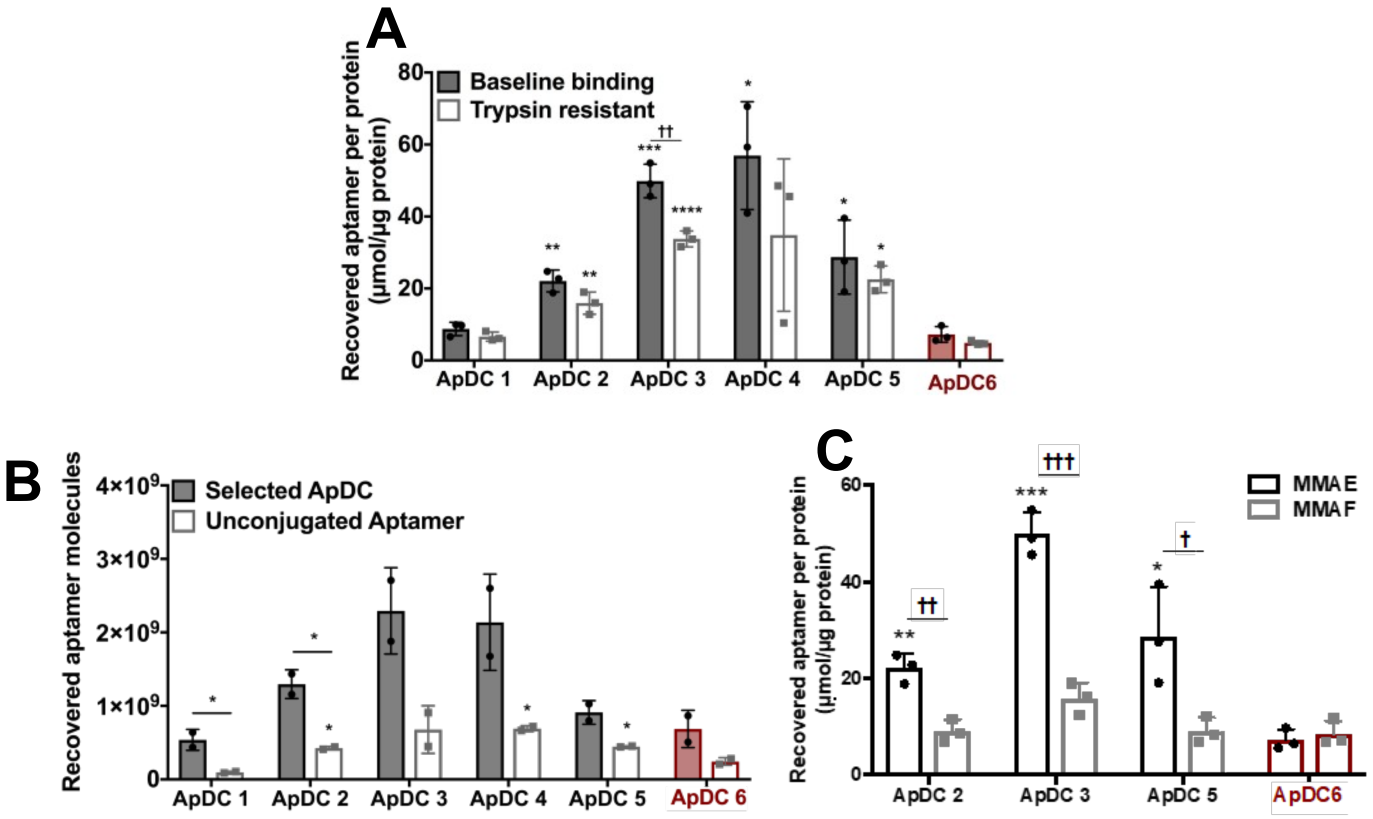


**Supplemental Figure S14.** A. ApDC binding to G39 cells after 30 min at 37°C before or after treatment with trypsin. Cells with were incubated with ApDC 72 h after seeding. B-C. ApDCs are evaluated for their target cell binding in cultures with and without MMAE (B) and MMAF substitution of MMAE (C).

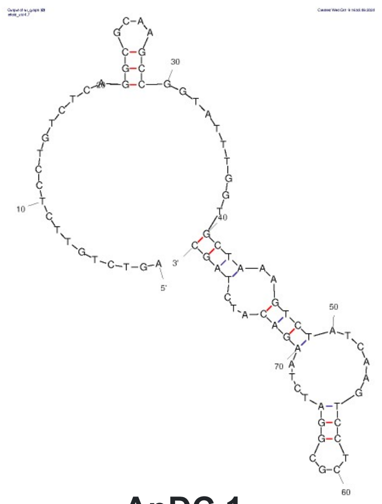

**ApDC 1**
**ΔG = -7.22 kcal/mol**

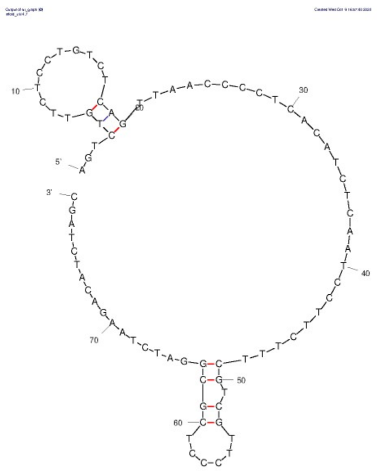

**ApDC 2**
**ΔG = -2.58 kcal/mol**

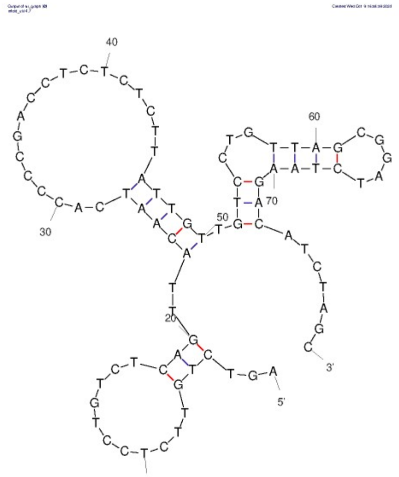

**ApDC 3**
**ΔG = -5.02 kcal/mol**

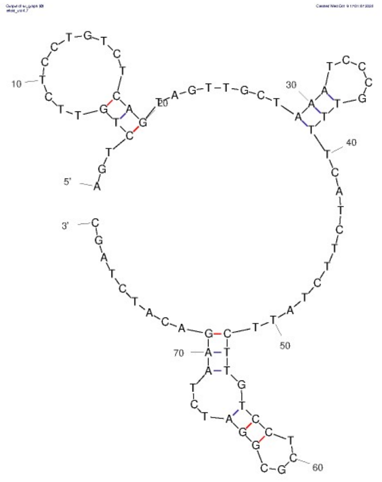

**ApDC 4**
**ΔG = -3.00 kcal/mol**

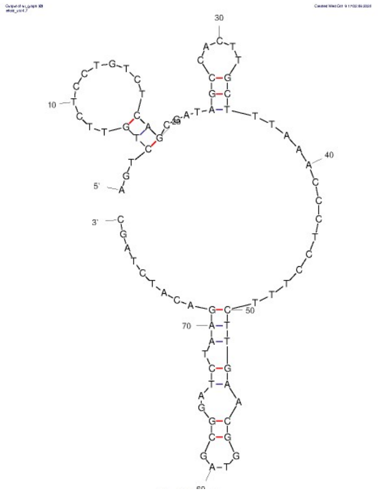

**ApDC 5**
**ΔG = -4.82 kcal/mol**

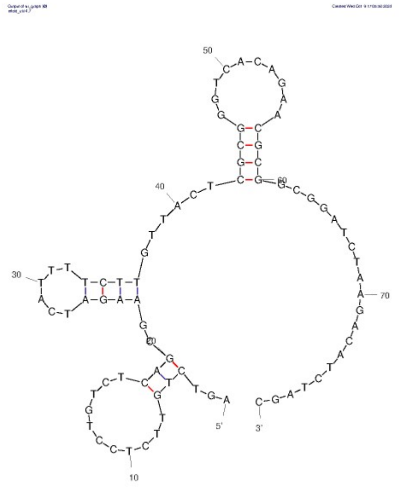

**ApDC 6**
**ΔG = -7.59 kcal/mol**

**Supplemental Figure S15.** Predicted secondary structures for the indicated aptamers as determined using mFold (Zuker M. *NAR*, 2003) (2).

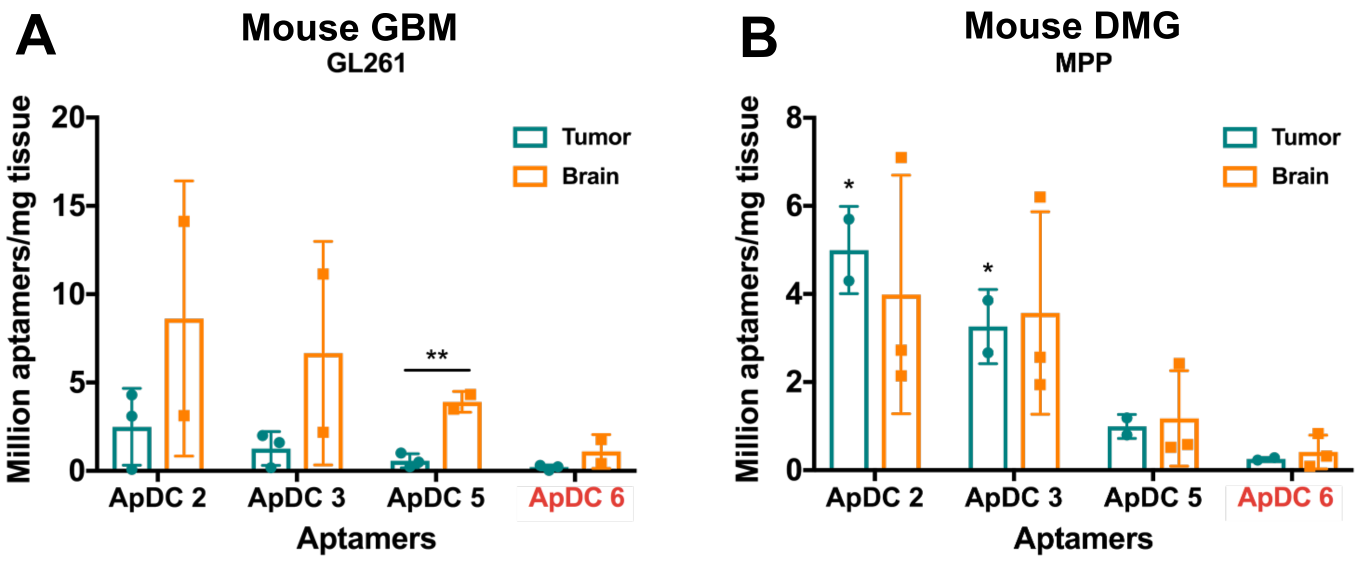


**Supplemental Figure S16.** A-B. Evaluating human G39 selected ApDC specificity to mouse high grade tumor models: GL261 (mouse GBM, A) and MPP (mouse DMG, B). H3K27MPP cells[3] were generously provided by Dr. Sameer Agnihotri's lab at the University of Pittsburgh.
